# Evaluating the impacts of microbial activity and species interactions on passive bioremediation of a coastal acid sulfate soil (CASS) ecosystem

**DOI:** 10.1101/2020.07.04.188110

**Authors:** Yu-Chen Ling, John W. Moreau

**Author notes:** Corresponding author: John W Moreau.

## Abstract

Microbial iron and sulfate reduction are the primary drivers of coastal acid sulfate soil (CASS) passive bioremediation schemes. Microbial sulfate reduction is the limiting step for pyrite formation, a desirable endpoint for CASS remediation. Little is known, however, about the impacts of microbial activity or species interaction on long-term iron and sulfur cycling in CASS ecosystems. Using a combination of molecular biology, geochemical speciation and artificial intelligence-powered computational modeling, we deduced from microbial activity patterns (RNA-based) and geochemical measurements a best-fit equation for predicting biogeochemical pyrite formation in a model CASS ecosystem. In addition to the time-dependent activities of key sulfate-reducing prokaryotic taxa (e.g. *Desulfobacteraceae*), this equation required methylotrophs (*Methylobacteriaceae*) and bacterial predators (*Bacteriovorax*) for best-fitting, suggesting that specific microbial interactions exert meaningful influences on CASS bioremediation efficiency. Our findings confirmed that CASS microorganisms act as an assemblage in response to rewetting by tidewater. Accurate predictions of long-term CASS bioremediation efficiency require modelling of complex and interdependent relationships between geochemical speciation and microbial activity.

**Highlights:**

- Coastal acid sulfate soil (CASS) is a global environmental issue.
- Microbial activity can be modelled quantitatively to predict CASS remediation.
- Sulfate-reducing prokaryotes (SRP) play a key role in CASS remediation.
- Predation on SRP with cultured representatives occurred during early wet-up.
- The above mechanism leads to increased activity among uncultured SRP.

## Introduction

In some coastal regions, human activity has accelerated the oxidation of naturally formed iron-sulfide minerals, triggering soil acidification and remobilization of heavy metals (Minh et al., 1997; Moreau et al., 2013; Sundström and Åström, 2002; White et al., 2007). These coastal acid sulfate soil (CASS) systems present an environmental problem worldwide (Ljung et al., 2009). More than $20 million CAD have been spent in the last century with an additional $7 million CAD system built in Halifax International Airport to control the acidic runoff (Hicks, 2003). There are about 3,000 hectares of farmland in Malaysia reported as acid sulfate soil, and ground magnesium limestone treatment costs around $1,130-3,820 USD per 10-hectare farm annually (Azman et al., 2014). The Australia Centre for International Agricultural Research spent around $0.85 million AUD on investigating acid sulfate soil remediation in Indonesia (project no. FIS/1997/022) (Myler, 2010). Australia is facing a $10 billion AUD “legacy” from acid sulfate soil (Fitzpatrick, 2003), even though Australian acid sulfate soil is only about 18% of the total 17 million hectare acid sulfate soils worldwide (Ljung et al., 2009).

Tidal inundation is one established strategy for remediating CASS (Powell and Martens, 2005), whereby flooding with seawater is used to neutralize acidity and induce (re)precipitation of metal-sulfides (Burton et al., 2007; Bush et al., 2004; Moreau et al., 2013). The driving force in this approach relies on the metabolic activity of anaerobic microorganisms that contribute up to 74% of the alkalinity in lime-assisted tidal treatment systems (Johnston et al., 2012). Anaerobic microbial “guilds”, such as iron- and sulfate-reducing prokaryotes (IRP and SRP, respectively) commonly use organic substrate to metabolise and generate Fe^2+^ and HS^-^. As Fe^2+^ and biogenic HS^-^ react, FeS forms as a poorly crystalline metastable phase (Meysman and Middelburg, 2005); conversion of this FeS to pyrite (FeS_2_) via S^0^ addition can then occur (Gagnon et al., 1995; Schoonen, 2004). The primary goal of CASS remediation is to neutralize acidity, and ultimately to re-precipitate iron sulfides (Burton et al., 2007; Bush et al., 2004), potentially as greigite or pyrite (Moreau et al., 2013), which are theoretically more stable phases (Billon, 2001).

The East Trinity wetlands (Cairns, Queensland, Australia) is a CASS system that experienced tidal inundation treatments since 2002 (Johnston et al., 2009b), which led to soil neutralization and heavy metal immobilization below the water table (Burton et al., 2011). In the East Trinity wetlands, microbial sulfate reduction only accounts for 7-13% of total alkalinity, compared to a 50-64% alkalinity increase from microbial iron reduction (Johnston et al., 2012). Previous studies have shown that biogenic HS^-^ is the limiting factor for pyrite formation in CASS ecosystems (Burton et al., 2011; Keene et al., 2010). Therefore, microbial sulfate reduction is the rate-determining step in CASS bioremediation.

In this study, we aimed to decrypt factors that influence microbial sulfate reduction in the East Trinity wetlands. SRP are not only the main driver of CASS bioremediation, but also play a vital role in the global carbon cycle. Furthermore, iron sulfide biominerals that precipitate as an indirect results of SRP activity can adsorb organic carbon and persist for years in anaerobic conditions (Picard et al., 2019). In our previous study, we found that environmental organic substrates are generally not a limiting factor for microbial sulfate reduction (Ling et al., 2015). However, microbial activity and species relative abundance have been shown to vary significantly in response to rewetting by tidewater (Ling et al., 2018). Therefore, we hypothesize that other microbial populations may impact SRP metabolic efficiency during different dry-down and wet-up (i.e. tidal) stages.

Previous studies have applied artificial intelligence methods to predicting: microbial diversity in soils (Park et al., 2011), net primary productivity in oceans (D’Alelio et al., 2020), cyanobacteria blooms in a freshwater lake (Tromas et al., 2017), nitrification processes in a sludge plant (Awolusi et al., 2016), and bioreactor performance in treating wastewater (Lesnik and Liu, 2017). Here, we test the significance of co-occurring environmental parameters and microbial network patterns in the East Trinity CASS system, and apply artificial intelligence modelling to predict the effects on pyrite formation. Similar to previous studies that focused on grassland microbial responses to rainfall or freshwater wetting leading to a peak in CO_2_ generation (Aanderud et al., 2015; Navarro-García et al., 2012; Placella et al., 2012), our study provides new data from a different type of environmental rewetting event (CASS tidal inundation) and coupled biogeochemical process (iron sulfide mineralization).

## 2. Materials and methods

### 2.1 Field site and soil sampling

The East Trinity wetland (145°80’E, 16°94’S) is located in a tropical estuary covered by mangrove trees and samphire (Hicks et al., 1999), across Chinaman Creek from Cairns, Queensland, Australia. The wetland was drained for sugarcane agriculture in the 1970’s, leading to severe soil acidification from enhanced oxidation of geogenic sulfide minerals (Powell and Martens, 2005). The practice of managed tidal inundation treatment was introduced in 2002 (Johnston et al., 2009b), and soil neutralization and heavy metals immobilization have since been reported for the soil layers below the water table (Burton et al., 2011).

Soil sampling and data preparation were performed using previously published methods (Ling et al., 2018). The sampling site consists of a 100-meter transect through the supra- to sub-tidal zones of East Trinity (Burton et al., 2011; Johnston et al., 2010; Ling et al., 2015). Duplicate sediment cores were collected in September 2013 across three sites (A: supra-, B: inter-, and C: sub-tidal zone, Fig. 1D) and four tidal stages (low tide: dry-down stage, flood tide: early wet-up stage, high tide: middle wet-up stage, ebb tide: late wet-up stage). These cores were then sectioned into seven intervals (1: 0-2 cm, 2: 2-4 cm, 3: 4-6 cm, 4: 6-8 cm, 5: 8-10 cm, 6: 12-14 cm, 7: 18-20 cm, Fig. 1D). One core from each duplicate set was used to assess microbial diversity and activity, and to quantitatively analyse environmental Fe and S speciation (Supplementary Table 1); the other core was used to evaluate reproducibility in molecular biology analyses. One biofilm sample was collected during the high tide stage from the surface water.

**Fig. 1.**
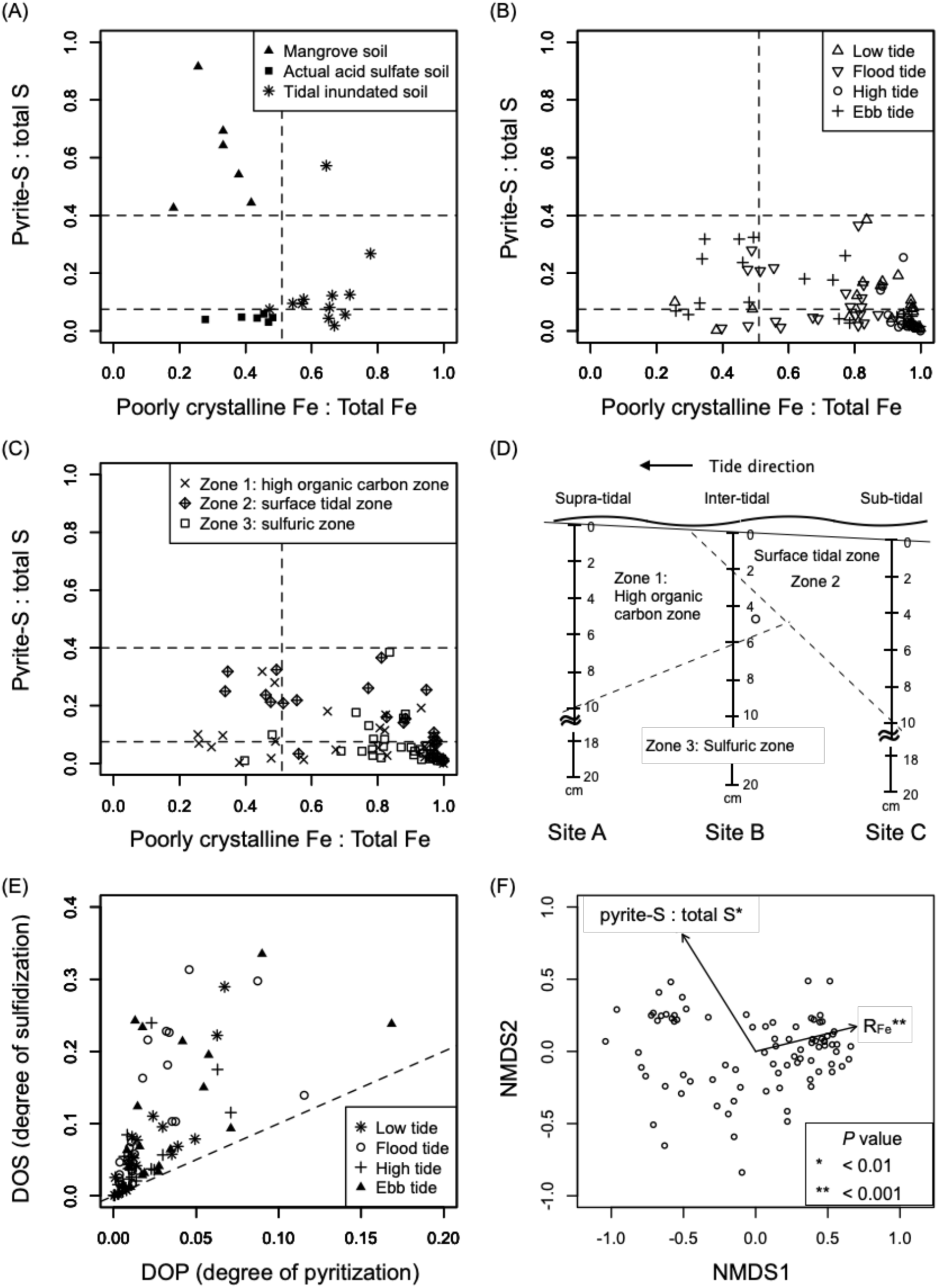
Field data of poorly crystalline Fe and pyrite-S fractions of (**A**) a previous study (Keene et al., 2010) and (**B, C**) this study. Samples were grouped based on tidal stages in (**B**), and on biogeochemical zones in (**C**). (**D**) The distribution of biogeochemical zones in the study site. (**E**) Correlations between DOS and DOP values in soil samples. (**F**) Beta diversity analysis comparing microbial taxonomic structural similarity of cDNA samples. The poorly crystalline Fe, pyrite-S, and organic carbon fractions were significantly correlated to microbial cDNA taxonomic structure.

In total, six different datasets were evaluated (Supplementary Table 1): microbial activity (based on RNA sequences), microbial distribution (based on DNA sequences), sulfate-reducing functionality (based on *dsrA* and *dsrB* genes from DNA and cDNA sequences), DNA and RNA dissimilarity (based on DNA and RNA sequences from one biofilm sample), protocol reproducibility (based on DNA sequences), and soil geochemistry (based on iron and sulfur speciation).

For checking the validity of sampling and bioinformatic protocols, duplicate soil subsamples from three sites, four tidal stages, and depths 2, 4, and 6, were selected to test for reproducibility (Supplementary Table 1). To prepare the datasets used for the evaluation of protocols, we extracted DNA from a total of 72 subsamples for 16S rRNA gene sequencing, and then analyzed the similarity of microbial taxonomic structures of these samples. If the sampling and bioinformatic protocols were reproducible, samples from the duplicate soil subsamples should share the highest similarity.

We selected triplicate soil subsamples from three sites, four tidal stages, and seven depths for microbial diversity and activity evaluation (Supplementary Table 1). The triplicate soil subsamples were homogenized before DNA and RNA extraction. RNA from a total of 84 samples were isolated to access microbial activity. For microbial populations, we homogenized subsamples across four tidal stages to get a total of 21 samples for DNA extractions. Because the amounts of samples were limited, DNA and cDNA (synthesized from RNA) samples from seven depths were homogenized for functional gene (*dsr*) sequencing, which contained 12 cDNA samples from three sites and four times, and DNA samples from three sites. Since soil DNA and RNA were not extracted from the same samples, we also extracted another DNA and RNA from a biofilm sample to evaluate the dissimilarity of microbial taxonomic structures between DNA and RNA sequences.

For DNA and RNA extractions, triplicate subsamples were collected from each core interval and preserved in 15 mL conical tubes with 2 mL of LifeGuard Soil Preservation Solution (MoBio Laboratory Inc., Carlsbad, CA, USA). The remaining soil samples from each interval were preserved in Petri dishes, and duplicate subsamples were selected afterward for geochemical speciation. All samples and subsamples were stored at −80 °C for subsequent analyses.

### 2.2 DNA and RNA extraction, 16S rRNA and dsr amplification

Total DNA and RNA from ∼1 g of soil were extracted following the PowerSoil DNA Isolation Kit protocol (MoBio Laboratory Inc.) and RNA PowerSoil Total RNA Isolation Kit protocol (MoBio Laboratory Inc.), respectively. Since our samples contained elevated organic C and salinity, we followed a troubleshooting protocol to use 1 g of soil, and incubated samples at ∼20°C in the nucleic acid precipitation step. DNA contamination in RNA samples was removed using the Ambion DNA-free kit (Ambion, Austin, TX, USA) or TURBO DNA-free kit (Ambion).

After RNA purification, cDNA was synthesized following the SuperScriptTM II Reverse Transcriptase (Invitrogen) protocol with random primers (Invitrogen) and the RNaseOUTTM (Invitrogen) reagent. We performed PCR (polymerase chain reaction) with three different primer sets to ensure there was (1) no detectable DNA amplification in the RNA samples, and (2) the DNA extraction and (3) cDNA synthesis were successful. The PCR using the following primers: 27F (5’-AGR GTT TGA TYM TGG CTC AG-3’)/518R (5’-ATT ACC GCG GCT GCT GG-3’), 926F (5’-AAA CTC AAA KGA ATT GAC GG-3’)/1392R (5’-ACG GGG GGT GWG TRC-3’) and U341F (5’-CCT ACG GGN GGC WGC AG-3’)/805R (5’-GAC TAC HVG GGT ATC TAA TCC-3’) (Klindworth et al., 2013). Dissimilatory sulfite reductase genes *dsrA* and *dsrB* were amplified from DNA and cDNA samples, with DSR1F+ (5’-ACSC ACT GGA AGC ACG GCG G-3’)/DSR-R (5’-GTG GMR CCG TGC AKR TTG G-3’) and DSRp2060F (5’-CAA CAT CGT YCA YAC CCA GGG -3’)/DSR4R (5’-GTG TAG CAG TTA CCG CA -3’) primer sets, respectively (Geets et al., 2006; Kondo et al., 2004; Perez-Jimenez et al., 2001).

Genomic DNA, genomic cDNA, *dsrA*, and *dsrB* samples were sent to Monash University Malaysia Genomic Facility for Illumina MiSeq sequencing (250-base pair, bp, 50 paired-end reads), with primer sets U341F/805 for genomic DNA and cDNA samples, DSR1F+/DSR-R for *dsrA* samples, and DSRp2060F/DSR4R for *dsrB* samples. Raw data were deposited in the Sequence Read Archive (SRA) of NCBI (PRJNA291811).

### 2.3 Bioinformatic analysis

We followed the MiSeq pipeline released by the MOTHUR authors (Kozich et al., 2013) to combine the forward and reverse sequences, trim low-quality bases and sequences with unexpected lengths, select unique sequences, cluster the unique sequences, detect chimera using UCHIME (Edgar et al., 2011), remove chimera, and classify the 16S and *dsr* sequences. Also, we used a phylotype-based approach for 16S OTU classification (Schloss and Westcott, 2011). After passing quality controls, sequences were aligned with the RDP database for classification. The 16S phylogenetic operational taxonomic units (OTUs) were defined based on taxonomic classification assigned by reference to the Ribosomal Database Project (RDP) (Cole et al., 2013) with an 80% confidence threshold and up to five mismatches using a naïve Bayesian classifier (Wang et al., 2007) (MOTHUR’s “classify.seqs” and “phylotype” commands).

Operational taxonomic identities were assigned to *dsr* sequences by reference to the published *dsrAB* databases (Müller et al., 2015), respectively, using a naïve Bayesian classifier (Wang et al., 2007) in MOTHUR. The *dsr* operational taxonomic units (OTUs) were defined by a distance of 0.1 (Müller et al., 2015). The most abundant sequence in each *dsrA* and *dsrB* OTU was selected as the representative sequence for building phylogenetic trees; 16S OTUs in cDNA and DNA datasets that shared the same family or genus names as *dsrA* and *dsrB* OTUs were selected, and the most abundant sequence in each OTU was selected for building the phylogenetic tree. All the steps were processed using MOTHUR.

### 2.4 Environmental parameter measurement

Soil moisture was measured by weighing mass loss in ∼5g soil after 24 hours of drying in an oven at 105°C. Measured values were used to normalise Fe and S speciation to per gram of dry soil.

Reduced inorganic sulfur species, including acid volatile sulfide (AVS), elemental sulfur (ES), and chromium-reducible sulfur (CRS), were extracted and measured sequentially following well-established protocols (Burton et al., 2011; 2009; 2008). AVS is defined as sulfur in iron monosulfide, ES is defined as elemental sulfur, and CRS is defined as sulfur in pyrite. AVS values can be converted to iron in iron monosulfide (FeS-Fe) using a 1:1 ratio, and CRS values can be converted to iron in pyrite (FeS_2_-Fe) using a 2:1 ratio (Table 2).

**Table 1.** The pyrite predicting equation and the taxonomic information of the selected taxa in the East Trinity coastal acid sulfate soil wetlands. Detailed taxonomic information of taxon089, taxon492, taxon694, and taxon160 are listed in Table S3.

| Formula | S ratio = 0.02406 + taxon0893 + 13.28*taxon1960 + 0.02406*taxon0941 + taxa0160*taxon0492 + taxon1730*taxon1739 + 1229*taxon0089*taxon0892*taxon1713 - taxon0694 - taxon1531*taxon1784 |  |  |  |  |  |  |
| --- | --- | --- | --- | --- | --- | --- | --- |
| Taxa | OTU | Kingdom | Phylum | Class | Order | Family | Genus |
| taxon0892 | OTU0035 | Archaea | Euryarchaeota | Thermoplasmata | unclassified | unclassified | unclassified |
| taxon0893 | OTU0036 | Archaea | Euryarchaeota | unclassified | unclassified | unclassified | unclassified |
| taxon0089 |  | Bacteria | Acidobacteria | Acidobacteria Gp3 |  |  |  |
| taxon0941 | OTU0084 | Bacteria | Actinobacteria | Actinobacteria | Acidimicrobiales | Acidimicrobiaceae | Ilumatobacter |
| taxon0492 |  | Bacteria | Actinobacteria | Actinobacteria | Actinomycetales | Microbacteriaceae |  |
| taxon1531 | OTU0674 | Bacteria | Proteobacteria | Alphaproteobacteria | Rhizobiales | Rhodobiaceae | unclassified |
| taxon0694 |  | Bacteria | Proteobacteria | Alphaproteobacteria | Rhizobiales | Methylobacteriaceae |  |
| taxon1713 | OTU0856 | Bacteria | Proteobacteria | Deltaproteobacteria | Bdellovibrionales | Bacteriovoracaceae | Bacteriovorax |
| taxon1730 | OTU0873 | Bacteria | Proteobacteria | Deltaproteobacteria | Desulfobacterales | Desulfobacteraceae | Desulfofaba |
| taxon1739 | OTU0882 | Bacteria | Proteobacteria | Deltaproteobacteria | Desulfobacterales | Desulfobacteraceae | unclassified |
| taxon1784 | OTU0927 | Bacteria | Proteobacteria | Deltaproteobacteria | Myxococcales | Polyangiaceae | unclassified |
| taxon0160 |  | Bacteria | Proteobacteria | Gammaproteobacteria |  |  |  |
| taxon1960 | OTU1103 | Bacteria | Proteobacteria | Gammaproteobacteria | Vibrionales | Vibrionaceae | Vibrio |

**Table 2.** The taxonomic information of bacterial predators in the East Trinity coastal acid sulfate soil wetlands.

| Kingdom | Phylum | Class | Order | Family | Genus |
| --- | --- | --- | --- | --- | --- |
| Bacteria | Proteobacteria | Deltaproteobacteria | Bdellovibrionales | Bacteriovoracaceae | Bacteriovorax |
|  |  |  |  |  | Peredibacter |
|  |  |  |  |  | unclassified |
|  |  |  |  | Bdellovibrionaceae | Bdellovibrio |
|  |  |  |  |  | Vampirovibrio |

AVS was extracted using 6 M HCl and two mL 1 M L-ascorbic acid, and captured using a NaOH-buffered zinc acetate solution, and then quantified by iodometric titration. ES was extracted from the AVS-extracted samples using toluene, and analyzed using high-performance liquid chromatography (HPLC) using a Dionex UltiMate 3000 system. CRS was extracted from the ES-extracted samples using acetone and ethanol, captured using a NaOH-buffered zinc acetate solution, and was quantified by iodometric titration.

Fe species included soluble Fe salts (MgCl_2_-Fe), crystalline Fe (CDB-Fe), and poorly crystalline Fe (the HCl-Fe minus FeS-Fe, Table 2); HCl-Fe, and CDB-Fe, were extracted and measured sequentially following established protocols (Claff et al., 2010), and MgCl_2_-Fe was extracted using 1 M MgCl_2_. The remaining samples were used to extract HCl-Fe using 1 M HCl; the amounts of extracted Fe(II) and total Fe were determined using spectrophotometry by the ferrozine method (Viollier et al., 2000). DCB-Fe was extracted from the residual samples using the dithionite-citrate buffer (DCB), and determined using inductively coupled plasma mass spectrometry (ICP-MS).

Soils were dried at 60°C for 48 hrs before crushing and analyzing for total Fe and S measurements at the Environmental Analysis Laboratory (EAL) of Southern Cross University (SCU). Total Fe was analyzed using inductively coupled plasma-mass spectrometry (ICP-MS) using a 1:3 HNO_3_/HCl digestion method, and total S was quantified using a LECO TruMAC CNS Analyser.

### 2.5 Data processing

We compared measured soil properties to results from previous studies of the same locations in East Trinity. Pyrite and poorly crystalline Fe fractions (defined as the ratios of pyrite-S: total S and poorly crystalline Fe: total Fe) in actual CASS, inundation-treated CASS, and undrained mangrove wetland from a previous study at the same site (Johnston et al., 2009c; Keene et al., 2010) were evaluated.

The degree of pyritization (DOP) and degree of sulfidization (DOS) provided indices to characterise depositional environments: DOS measures the extent of reactive iron transformed into iron sulfide minerals (Boesen and Postma, 1988), while DOP represents reactive iron transformed to pyrite (Berner, 1970). In this study, we calculated the difference between DOS and DOP to deduce the extent of reactive iron transformation to iron monosulfide, but which was not yet converted to pyrite. DOS and DOP were calculated as follows (Berner, 1970; Boesen and Postma, 1988):

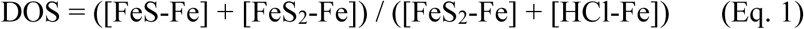

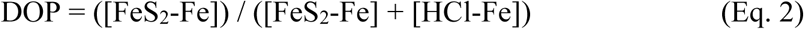

where FeS is iron monosulfide, and FeS_2_ is pyrite.

Sequences were rarefied before calculating similarities among samples using the “dist.shared” and “sub.sample” commands in MOTHUR. The “dist.shared” command generated a distance matrix for dissimilarity analyses. The “sub.sample” command normalized samples to the same size by selecting sequences randomly prior distance calculating. Sample normalization and distance calculation were repeated for 1,000 times and generated an average result of distance matrix for subsequent dissimilarity analyses using MOTHUR. The taxonomic similarity between cDNA and DNA (16S rRNA gene based) in soil and biofilm samples was calculated using the Jaccard index (Jaccard, 1908) and visualized by non-metric multidimensional scaling (NMDS) using MOTHUR software. The test statistic and associated significance value (ANOSIM) (Clarke, 1993) of these four datasets were calculated using MOTHUR.

To evaluate whether pyrite and poorly crystalline Fe ratios drove observed differences in cDNA taxonomic-based microbial community structures, we first calculated taxonomic dissimilarity between 16S OTUs in cDNA rarefied samples using the Bray-Curtis index, then we tested these environmental factors for correlation, with significant correlations (*p* < 0.05) shown as vectors on the non-metric multidimensional scaling (NMDS) using the *R* package vegan.

We used the artificial intelligence package Eureqa (Schmidt and Lipson, 2013; 2009) to find a best-fit equation for predicting soil pyrite ratios from microbial cDNA (16S based) contents (i.e., to construct a “pyrite predicting equation”). Eureqa first creates random equations based on the provided dataset, then evaluates the fitness of every equation. These preliminary best-fit equations are selected and are used to generate a new round of searching. In the new round of searching, new equations are formed by varying the previous equations and retaining the equations that better fit the dataset. The 84 soil samples were ranked based on pyrite-S fractions, and then divided into two sets (Fig. S4A). One set was used to train Eureqa to find the best-fit equation, and the other set was used to validate results (Fig. S4A). Taxa present in less than 25% of the samples were removed from both the training and validation sets. The accumulated relative abundances of selected OTUs from each sample were calculated to evaluate that filtering criterion.

To assess potential environmental and microbial parameters controlling SRP activity, geochemical data (e.g., concentrations of pyrite, iron monosulfide, or organic carbon) and variations in cDNA levels of selected taxa and metabolic guilds (SRP, *Bdellovibrionales, Acidobacteria GP 3*, and *Gammaproteobacteria*) in different dry-down and wet-up stages were quantitatively compared.

To evaluate the response of microbial interactions to rewetting by tidewater, all 84 cDNA samples were divided into four groups based on collection timepoints. Each group contained 21 subsamples across depths and tidal zones used to perform microbial network analysis, in order to investigate microbial interaction patterns across the study site at specific dry-down and wet-up stages. OTUs present in less than 50% of samples from each tidal stage were removed from the cDNA datasets. Microbial interaction networks were inferred using CoNet (Faust et al., 2012) and visualized using Cytoscape (Shannon et al., 2003). Correlation between microbial cDNA abundances were measured using the Pearson and Spearman correlations, Bray–Curtis distance and Kullback–Leibler divergence measurement, then the best predictions were selected. A well-established approach was used for multiple-testing correction (Benjamini and Hochberg, 1995), and p-values were merged using the method of Brown (Brown, 1975). Only the relationships that displayed significant correlations (p < 0.05) were visualized as networks. Microbial networks were reconstructed, containing OTUs that were present in more than 50% of samples across all four tidal stages, and displayed significant correlations with each other. In these networks, each node represents one OTU, while connections between two nodes represent an interrelationship between two OTUs. Due to the complexity of node relationships, sub-networks were constructed to focus on potential predator-prey relationships. Only the first and second neighbors of potential predators were kept in these sub-networks.

### 2.6 Plotting and data processing software

The software R (Team, 2013) with packages gplots (Warnes et al., 2009), MASS (Venables and Ripley, 2002), vegan (Oksanen et al., 2015), and plyr (Wickham, 2011) were used for statistical analysis and data visualisation. We also used the AI software package Eureqa to analyse datasets for network correlations. The software packages TreeGraph 2 (Stöver and Müller, 2010), Cytoscape (Shannon et al., 2003) with application CoNet (Faust and Raes, 2016), iWork Keynote and Office Excel were also used to visualise data and generate plots.

## 3. Results

### 3.1 Replication evaluation and overall microbial taxonomic structures

Our previous work showed that samples from the same soil depth interval displayed the highest similarity of microbial DNA taxonomy-based microbial community structure, confirming our sampling reproducibility (Ling et al., 2018). We identified 1170 OTUs from a total 107 16S sequencing samples for microbial distribution and activity, DNA and RNA dissimilarity evaluations. A significant separation between soil DNA and soil RNA taxonomic structure (16S based) was detected by ordination NMDS (Fig. S1A); ANOSIM analysis displayed an *R*-value of 0.45, *p* < 0.001 (Fig. S1B). The R values are between 1 and −1, in which positive numbers suggest that there are more similarities within groups than between groups; negative numbers indicate vice versa.

### 3.2 Geochemical characteristics

The chemical profiles of iron sulfide minerals show that site C (subtidal zone) contains the highest amount of iron monosulfide and pyrite among the three sampling sites, and of iron monosulfide accumulated in the flood and ebb tides (Fig. S2). By comparing the poorly crystalline Fe and pyrite-S contents to values from a previous study (Fig. 1A), we could semi-quantitatively assess the degree of bioremediation progress achieved to date in the East Trinity wetland site. None of our samples contained the pyrite-S: total S ratios representative of “undisturbed” mangrove soils (Fig. 1B). Instead, our soil samples displayed variable levels of poorly crystalline Fe and pyrite-S. Generally, samples that contained lower pyrite and poorly crystalline Fe levels originated mostly from the organic-enriched zone (Figs. 1C and 1D), whereas samples that contained higher pyrite-S: total S ratios were collected during ebb tide conditions (Fig. 1C). DOS values were higher than the DOP values in all of our samples (Fig. 1E), suggesting that Fe was present in the more reactive iron monosulfide phase instead as pyrite. Variations of cDNA taxonomic structure were correlated with soil pyrite and poorly crystalline Fe ratios (Fig. 1F).

### 3.3 Prediction of soil pyrite ratios

We selected 326 and disregarded 686 OTUs (from a 16S cDNA-constructed database) for predicting soil pyrite ratios, the selected OTUs constituted more than 98% of relative abundances (Fig. S3). The best-fit pyrite equation, based on 72 samples in the training set, displayed a Pearson coefficient of *r* = 0.92 with *p* < 2.2e-16 (Fig. S4B). This equation predicted soil pyrite ratios well for the validation dataset (*r* = 0.88 with *p* < 0.001, Fig. S4C). The equation contained two archaeal taxa (*Euryarchaeota*), 11 bacterial taxa (*Acidobacteria, Actinobacteria*, and *Proteobacteria* classes *Alpha*-, *Delta-*, and *Gamma*-*proteobacteria*) (Table 1). Among the selected taxa, we identified taxa 1730 and 1739 as SRP, taxon0694 as a methylotroph (Kuever, 2014), and taxon1713 as a bacterial predator (Table 1). We also observed taxa that could be assigned to early and delayed responders identified from different wet-up stages in our previous study (Ling et al., 2018). *Acidobacteria GP 3* displayed a higher cDNA contents during flood and high tides (taxon0089, Fig. S4D), and taxa within the *Gammaproteobacteria* had the highest cDNA contents during the ebb tide (e.g. taxon0160, Fig. S4F).

### 3.4 Microbial activities and chemical variations during different tidal stages

Seven SRP were classified at the family or genus level from *dsrA* and *dsrB* sequence datasets, with six of these also identified in the 16S dataset (Fig. S5). One SRP taxon found with 16S, but not *dsrA* or *dsrB*, sequences was of order *Desulfuromonadales*. In the following analysis, classified SRP were referred to as *dsrA* and *dsrB* targeted OTUs (Fig. S5) and *Desulfuromonadales*. Ten of the SRP taxa identified using *dsrA* and *dsrB* genes are presently uncultured (Fig. S5). The number of uncultured SRP able to be identified was higher when using *dsrB*, as compared to *dsrA*, genes (Fig. S5). Thus we represented cDNA variability of uncultured SRP across tidal stages using reverse-transcribed *dsrB* cDNA sequences (Fig. 2E). These uncultured SRP displayed increased *dsrB* cDNA during the middle and late wet-up stages (high and ebb tides), in contrast to *dsrB* cDNA from flood and high/ebb tides (*t*-*test, P* < 0.05, Fig. 2E). In contrast, SRP OTUs most closely related to cultured representatives increased in cDNA during the early and middle wet-up stages (flood and high tides), and decreased during the late wet-up stage (ebb tide). Average cDNA values during ebb tide showed significant differences from the other three tidal stages (*t*-*test, P* < 0.05, Fig. 2D).

**Fig. 2.**
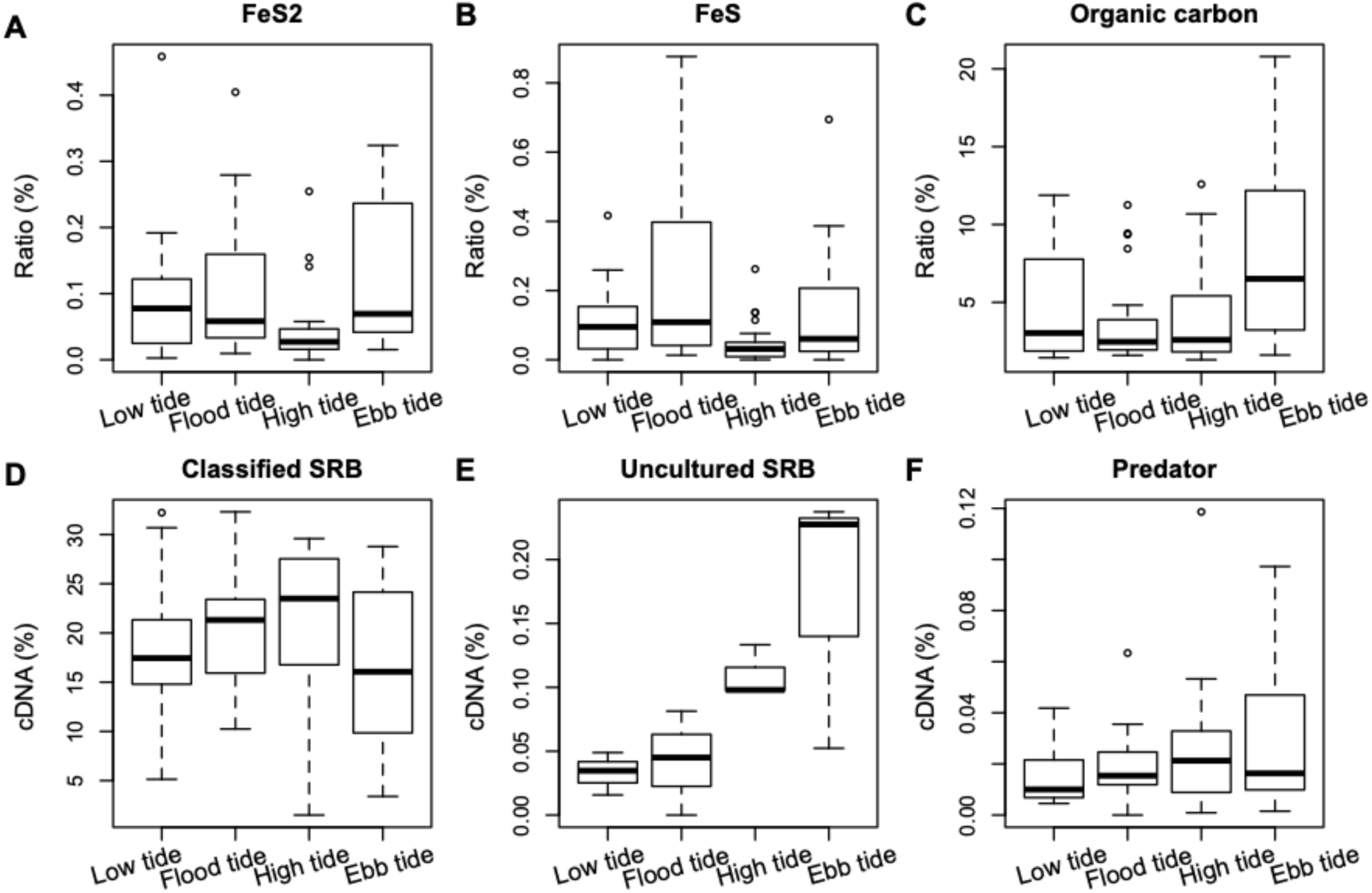
Environmental parameters and microbial activities, including (**A**) pyrite contents, (**B**) iron monosulfide content, (**C**) organic carbon content, (**D**) classified SRP activity, (**E**) uncultured SRP activity, and (**F**) predator activity across different dry-down and wet-up tidal stages. In the boxplots, upper and lower bars represent maxima and minima, respectively, excluding outliers plotted as points. The upper and lower portions of boxes from the first and third quartiles, respectively, while the lines inside boxes are sample medians.

To interpret changes in cDNA levels for all SRP taxa, we co-analyzed (1) changes in environmental parameters, and (2) microbial predator abundance over different dry-down and wet-up tidal stages. Pyrite and iron monosulfide accumulated at flood and ebb tides (Figs. 2A and 2B), and organic carbon content was highest at ebb tide (Fig. 2C). Five predator genera were identified within order *Bdellovibrionales*, including *Bacteriovoracaceae* and *Bdellovibrionaceae* families (Table 2), two well-known predatory microorganisms (Pineiro et al., 2007). The cDNA levels of microbial predators displayed similar patterns to known SRP, with increasing cDNA levels during flood and high tides, and decreasing cDNA at ebb tide (Fig. 2F). The cDNA levels of known SRP and predators were proportional only at flood tide (Figs. 3A-D).

**Fig. 3.**
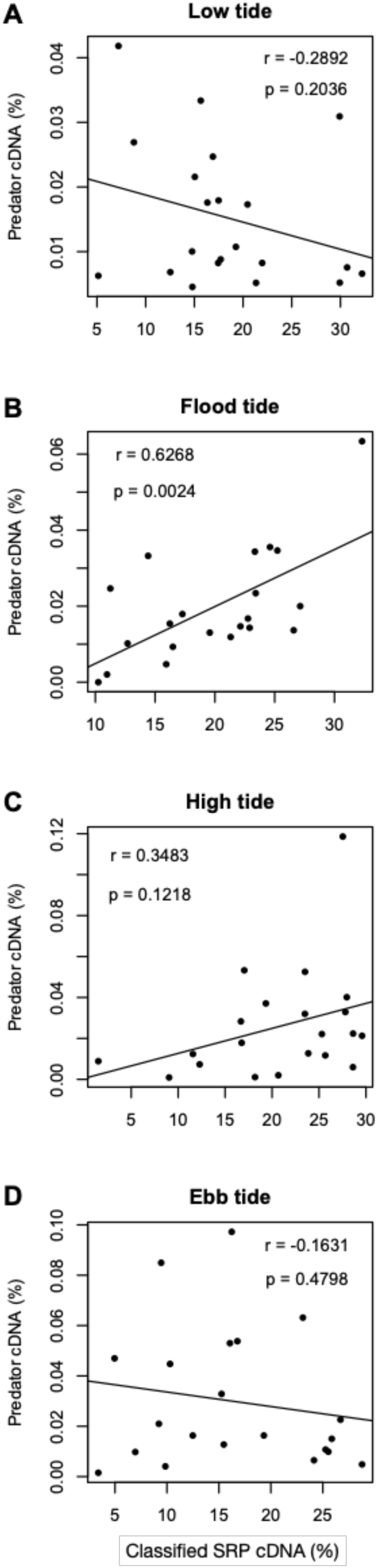
Correlations between cDNA contents of classified SRP and predators in (**A**) low tide, (**B**) flood tide, (**C**) high tide, and (**D**) ebb tide.

The numbers of predators’ first neighbors in microbial networks were 1, 16, and 9, at low and ebb tides, flood tide, and high tide, respectively (Fig. 4). Among these first neighbors, 2 and 1 known SRP taxa negatively correlated with predator taxa during the flood and high tides, respectively. OTU 1034 was first neighbor to predators taxa *Bacteriovoracaceae* and *Bdellovibrionaceae* at both ebb and high tides, and is most closely related to genus *Thioprofundum*, known to fix CO_2_ and oxidize sulfur (Mori et al., 2011; Watts et al., 2017). OTU 809, most closely related to genus *Burkholderia* (Estrada-de los Santos et al., 2001), capable of fixing nitrogen, was first neighbor to predator taxa at low tide.

**Fig. 4.**
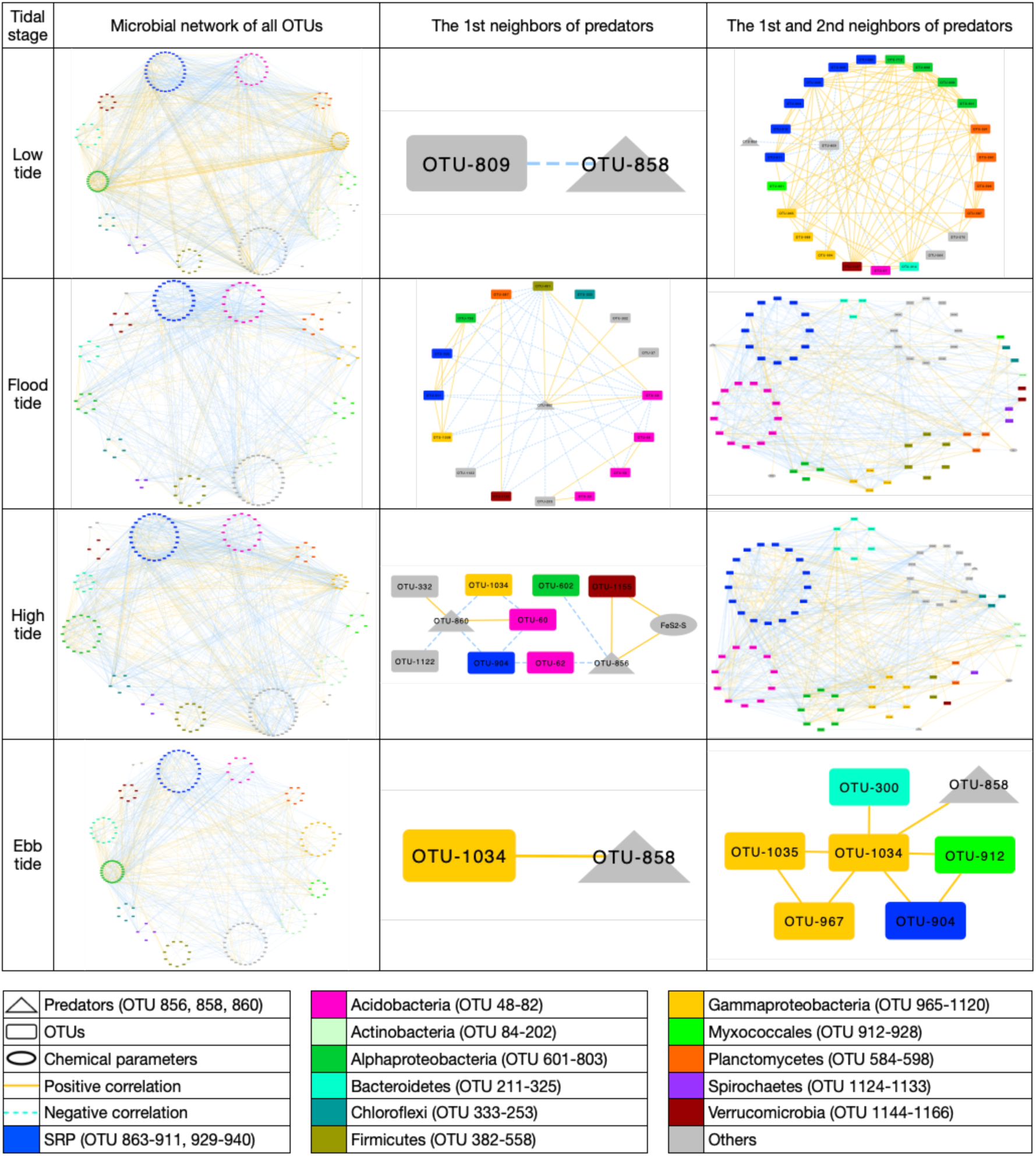
Microbial networks of all OTUs, predators’ first and second neighbors across four tidal stages. Each node represents an OTU or environmental parameter, and edges that connect OTUs or parameters represent either positive (yellow line) or negative (blue dashed line) correlations between the two nodes.

In microbial network analyses, OTUs needed to be present in more than 50% of samples from each tidal stage, and to display a significant correlation to other OTUs, in order to be visualized as networks. OTUs that were present in more than 50% of samples from each tidal stage were: 244, 153, 174, 243, at low, flood, high, and ebb tides, respectively. The relationships between OTUs with *p*-values above the threshold (0.05) were then removed. In the end, the total number of nodes (OTUs) were 200, 138, 163, and 178 at low, flood, high, and ebb tides, respectively (Fig. 4). There were 1231, 638, 912, and 659 connections among nodes for low, flood, high, and ebb tide networks, respectively. Predators’ first and second neighbor relationships during flood and high tides outnumbered those of other tidal stages (Fig. 4). The numbers of predators’ first and second neighbors in microbial networks were 24 at low tide, 67 at flood tide, 75 at high tide, and 7 at ebb tide. Known SRP were among the first and second neighbors to predator taxa across all four tidal stages.

## Discussion

Soil microorganisms can contain a higher abundance of inactive microbes than other environments, such as freshwater, activated sludge, and the human gut (Lennon and Jones, 2011). These inactive microorganisms have low metabolic rates when environmental conditions become unfavourable, and decoupling occurs between active and total microbial populations (Jones and Lennon, 2010). A previous study observed significant separation between standing (DNA-based) and active (RNA-based) communities in soil systems (DeAngelis et al., 2010), which was also found in our study site (Fig. S1). To observe relationships between environmental and microbial dynamics in a more rapid period of land transition, researchers have suggested using RNA-based community structure rather than DNA (Meyer et al., 2019). Tidal inundation treatment imposes rapid environmental changes on coastal microorganisms, and therefore we used cDNA (RNA-based) communities to link with soil mineralization for this study.

Because rRNA (cDNA) content may not always be directly proportional to growth rate, and some microorganisms can contain more rRNA in dormancy (Blazewicz et al., 2013), we used the variation in cDNA relative abundances as an indicator of microbial activity. Since growth is only part of cellular activity (Hoehler, 2004), the cDNA variability to which we refer represents overall activity and not solely growth. Variation in cDNA content can be viewed as a change of total activity (Ling et al., 2018), because RNA synthesis costs energy (Wagner, 2005); thus we treat increasing cDNA levels as increasing activity, and vice versa. The following analyses of microbial activity (Figs. 2D-F, 3) relied on changing cDNA contents across different dry-down and wet-up tidal stages, and the constructed pyrite-prediction equation was based on cDNA variations across 84 samples (Figs. S4A-C). The microbial interaction networks (Fig. 4) were built by comparing the cDNA variability of OTUs during each dry-down or wet-up stage.

### 4.1 Passive bioremediation progress in the East Trinity wetland

A previous study compared the compositions of soil iron and sulfide compounds in soils from three different bioremediation stages: mangrove soils (represented as undisturbed soils), actual acid sulfate soils, and tidally inundated CASS soils (Keene et al., 2010). The mangrove soils contained high amounts of pyrite (Fig. 1A) that was oxidized during drainage to form Fe(III) minerals; reductive dissolution of Fe(III) minerals during tidal inundation leads to accumulation of poorly crystalline Fe and re-precipitation of pyrite (Fig. 1A). The soil samples in this study contained different degrees of soil pyrite and poorly crystalline Fe ratios providing a range of mineral phase and potential transformations. A small number of samples displayed low pyrite and poorly crystalline Fe ratios (Figs. 1B and 1C), characteristic of actual acid sulfate soils. These samples were located in the high organic carbon zone (Figs. 1C and 1D), where soil pH was below 5.5 (Ling et al., 2018; 2015), conditions which are consistent with the formation of CASS. Soils in the supra-tidal zone were exposed to air for a longer time than those in inter- and sub-tidal zones, causing pyrite oxidation and release of acidity. Samples containing the highest pyrite ratios and lowest poorly crystalline Fe ratios were collected during the flood and ebb tide periods (Fig. 1B), confirming that tidal inundation provides the primary CASS remediation mechanism. Previous studies reported that soil pH in the East Trinity wetland increased from ∼3 to ∼6 after five years of tidal inundation treatment (Hicks et al., 1999; Johnston et al., 2009a).

Soil pyrite and poorly crystalline Fe ratios in our samples also suggested that remediation is ongoing, and has not yet resulted in pyrite contents as high as those associated with undisturbed mangrove soils. Relatively high DOS and low DOP values in our samples (Fig. 1E) suggested that iron monosulfides, instead of pyrite, comprised the bulk of iron sulfide mineral phases in our system. Many samples contained abundant reactive iron (Fig. 1B), favoring iron monosulfide accumulation and limiting the transformation of AVS and ES to pyrite (Keene et al., 2010). This observation is consistent with results from a previous study of the same site, in which sulfide concentrations were below detection limits (Burton et al., 2011). Compared to pyrite that is more stable during tidal inundation stages, soils contained higher amounts of iron monosulfide during the flood and ebb tides, (Fig. S2). This finding supports the previous study which stated that pyrite is considered to be a stable compound (Billon, 2001). Iron monosulfide, however, could form black ooze with organic matter and cause rapid de-oxygenation when mixed with water (Bush et al., 2004).

### 4.2 Microbial activities controls pyrite formation

The taxonomic structure of active microbial populations correlated with soil pyrite and iron ratios (Fig. 1F), reflecting concomitant changes in microbial activities and soil iron and sulfur cycling. The pyrite prediction equation accurately predicted soil pyrite ratios based on cDNA content (Fig. S4C), confirming that microbial activity mainly controlled pyrite formation.

The best-fit pyrite prediction equation contains multiple metabolic guilds, such as SRP, methylotrophs, and predators (Table 3), although clearly SRP comprise the dominant metabolic guild in the CASS system (Ling et al., 2018; 2015), and their metabolic product (sulfide) is the limiting factor to forming pyrite. A previous study reported that microbial predation can decrease SRP growth and metabolic yields (Qiu et al., 2016). *Methylobacteriaceae* is one of the most abundant taxa in the sulfide-bearing sediments that experience periodic dry-down and wet-up (Jones et al., 2017), and is also part of the best-fit pyrite prediction equation. Thus, the accuracy of this relationship was improved by inclusion of potential microbial network interactions.

In our previous study, we observed sequential microbial guild resuscitations across different dry-down and wet-up tidal stages (Ling et al., 2018). So called “early responders” increased in activity during the initial stages of seawater inundation, while “delayed responders” increased in activity during late wet-up stage. The pyrite prediction equation also accounts for such time-dependence in the activity of different microbial taxa (Figs. S4D and S4E). For example, *Acidobacteria Gp 3* (taxon0089) and *Gammaproteobacteria* (taxon0160) were identified as early and delayed responders, respectively, in our previous study (Ling et al., 2018). Sequential microbial resuscitations might account for the observed pyrite and iron monosulfide accumulations during two distinct periods (early and late wet-up stages, Figs. 2A and 2B). Since microbial activity changes quickly (over hours) during tidal inundation treatment (Ling et al., 2018), it is possible that more than one metabolic group was ultimately responsible for net iron sulfide formation.

We note that the best-fit pyrite prediction equation is based on four different dry-down and wet-up tidal stages (symmetrically across two days to avoid night sampling in crocodile inhabited waters). Some factors that could influence bioremediation efficiency over a more extended time period were: minerals preserved in organic matter, increasing temperature, changing rainfall patterns, and day/night (i.e. wetting/drying) shifts. We observed that iron-sulfide minerals preserved labile organic carbon from microbial degradation in the system (Ling et al., 2015). Forming iron sulfide minerals is the primary goal to remediate soil acidity in the CASS systems, and higher ratios of iron sulfide minerals could change organic matter availability and thus further influence microbial metabolic efficiency. In the future, a greater contrast in precipitation between wet and dry seasons is expected due to climate change (Trenberth *et al*., 2014). It is predicted that the temperature will be warmer by 1.2 to 4.2°C, and the rainfall will change by −40 to +5% in the southeast region of Australia by 2090 (Reid and Mosley, 2016). When the evaporation rates exceed precipitation rates due to higher temperature, such as with summertime in the Netherlands, coastal acid sulfate soils will release more sulfuric acid to nearby ecosystems than during any other season (Vermaat *et al.*, 2016). Microbes have displayed resuscitation sequencing in response to rewetting events such as rainfall (Placella et al., 2012) and tidal inundation (Ling et al., 2018). Therefore, prolonged rainfall may generate different microbial metabolic yields to multiple short periods of rain. A previous study showed that phagotrophic protists are more active at night, compared to daytime, in a subtropical gyre (Hu et al., 2018), while another study reported that the existence of protists increases microbial sulfate reduction activity in a sludge reactor (Hirakata et al., 2016). It is possible that protists engulfing prokaryotes displayed different activity patterns between day and night, hence impacting the overall dynamics of bioremediation. For long-term remediation efficiency in acid sulfate soils, such environmental factors should be considered and studied in more detail.

Compared to soil models such as RothC and CENTURY that are designed for predicting soil organic carbon (Farage et al., 2007; Molina et al., 2017), artificial intelligence packages provide flexible options for researchers to customize modelling targets. For example, previous studies have used Eureqa to predict bacterial assemblages in marine systems (Larsen et al., 2012), and microbial taxonomic and functional structures in acid mine drainage (Kuang et al., 2016). Here we have modelled microbial activity to predict soil pyrite ratios in coastal acid sulfate soils. However, results from artificial intelligence packages are not always straightforward and often require further interpretation to decrypt key relationships between the parameters that are listed in the predictive algorithms.

### 4.3 Microbial interactions changed during different dry-down and wet-up tidal stages

Although organic carbon fuels bacterial sulfide generation, we did not observe variations in organic carbon to correlate with pyrite or iron monosulfide levels (Figs. 2A-C). Instead, when seawater entered the CASS system during flood tide, we observed that cDNA contents of SRP increased and were accompanied by pyrite and iron monosulfide accumulations. However, SRP decreased their cDNA content at ebb tide, whereas pyrite and iron monosulfide levels exhibited another concentration peak at this stage, obfuscating this relationship somewhat. Furthermore, SRP decreased in activity at a tidal stage during which their environment still contained a high abundance of organic carbon, suggesting that other parameters controlled SRP activity.

The identification of classified and uncultured SRP (Fig. S5) allowed us to compare physiological characteristics between these two groups. Order *Desulfuromonadales* was observed in 16S datasets only, but in neither *dsrA* or *dsrB* sequences, which suggests that *Desulfuromonadales dsr* genes may differ from those of strain *Desulfuromonadales* bacterium Tc37 (reference sequence). Compared with classified SRP that increased in cDNA content during flood and high tides (Fig. 2D), uncultured SRP increased in cDNA content quickly during ebb tide (Fig. 2E). Both classified and uncultured SRP utilize sulfate as the electron acceptor, and ebb tides displayed the highest organic carbon contents during different dry-down and wet-up tidal stages (Fig. 2C), indicating that the decreasing activity of classified SRP during the ebb tide was not limited by the availability of organic substrate. Both predators and classified SRP increased their cDNA levels during flood and high tides (Fig. 2F), and corresponding changes in predators and classified SRP cDNA levels were observed at flood tide (Figs. 3). This observation supports a “kill the winner” hypothesis (Maslov and Sneppen, 2016), in which SRP forming dominant metabolic guild in the system become substrates for microbial predation.

To further investigate microbial predator-prey relationships, we built microbial association networks for each tidal stage (Fig. 5). The interactions of predators during different dry-down and wet-up tidal stages showed that they played a “keystone” role during early and middle wet-up stages (flood and high tides). Keystone species have significant influence on the community, and form higher level nodes in microbial co-occurrence networks (Berry and Widder, 2014). We also observed that keystone species consisted of the predators’ first neighbors in microbial networks during early and middle wet-up stages (flood and high tides). Classified SRP are among the predators’ first and second neighbors, which supports our hypothesis that predation influenced classified SRP activity during these wet-up stages. Microbial predation on classified SRP likely created opportunities for uncultured SRP, potentially having a narrower environmental niche, to become more active during delayed wet-up stage (ebb tides). Since classified SRP taxa decreased in activity during ebb tides, while pyrite and iron monosulfide still accumulated, we may assume that uncultured SRP also contributed significantly to total iron sulfide formation.

**Fig. 5.**
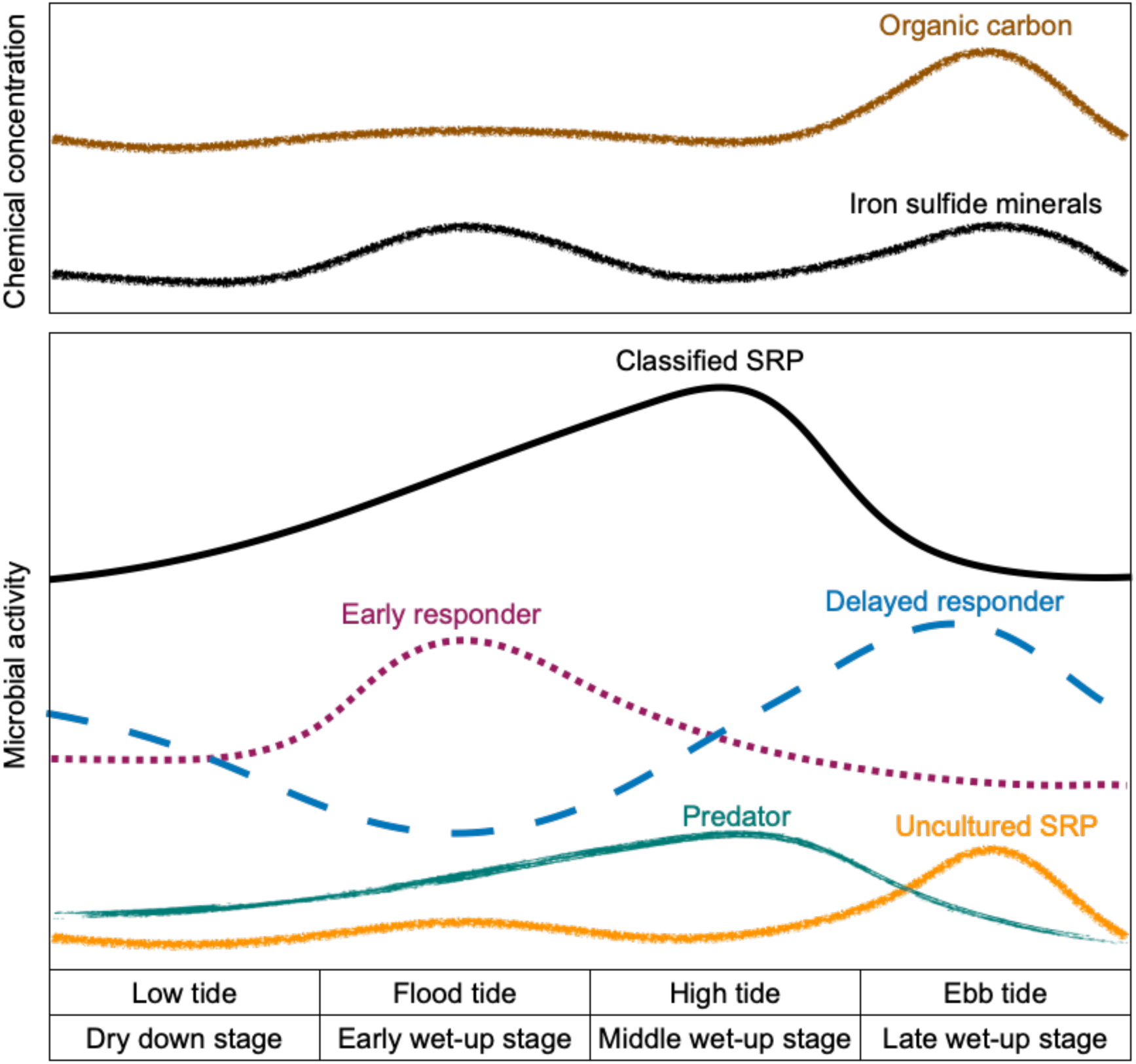
Conceptual model showing the trend in relative activity of classified and uncultured SRP, predators, early and delayed responders over different dry-down and wet-up tidal stages.

In summary, our observations confirm that microbial activity controls iron and sulfur speciation, and serves as an accurate predictor for pyrite (re)precipitation in CASS bioremediation. We discovered that microorganisms displayed at least two different interaction patterns that influenced iron sulfide formation at the early and late wet-up stages (Fig 5). Classified SRP were early responders to tidal inundation and became abundant taxa in the system, which induced microbial predation and decreased their activity during late wet-up. Microbial predation on classified SRP taxa created ecological space for uncultured SRP taxa to proliferate and further drive iron sulfide formation. Uncultured SRP increased their activity quickly during ebb tides, and also contributed to net pyrite formation. Our work reveals the need to consider a complex interplay of geochemical parameters and microbial interactions in the (bio)remediation of CASS ecosystems.

## Acknowledgements

The authors acknowledge funding to JWM from the Australian Research Council (ARC LP110100732). We thank Drs. Kathryn Holt, Shu Mei Teo and Andre Mu (University of Melbourne) for access to and assistance with computational biology resources, Dr. Caitlin M. Gionfriddo (Oak Ridge National Laboratory) for important advice on writing and manuscript structures, and Mr. Steve Wilbraham (Queensland Government) for invaluable assistance with fieldwork and sampling at the East Trinity field site.

