## Supplementary material for "Evaluating the impacts of microbial activity and species interactions on passive bioremediation of a coastal acid sulfate soil (CASS) ecosystem"

### 1. Supplementary Figures

**A**

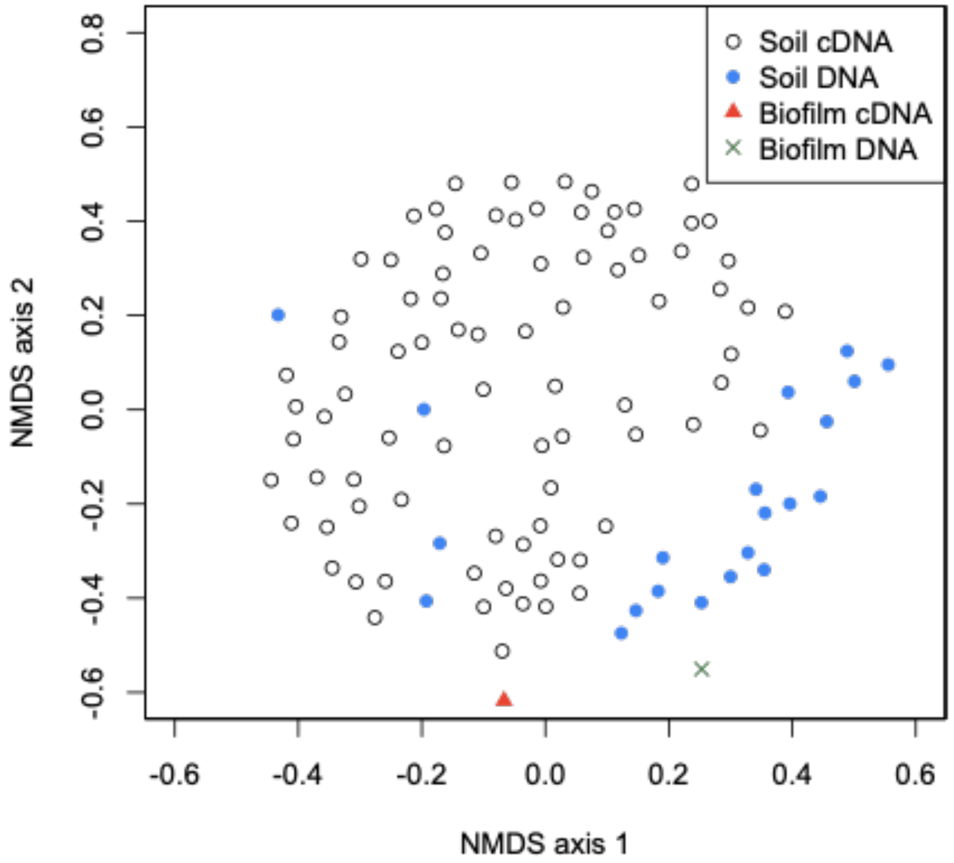


**B**

| ANOSIM comparison | R-value | P-value |
|---|---|---|
| ● Soil DNA vs. ○ soil cDNA | 0.447212 | <0.001 |
| ✕ Biofilm DNA vs. ▲ biofilm cDNA | nan | <0.001 |
| ● Soil DNA vs. ✕ biofilm DNA | 0.556463 | 0.049 |
| ○ Soil cDNA vs. ▲ biofilm cDNA | 0.70595 | 0.01 |
| ● Soil DNA vs. ▲ biofilm cDNA | 0.697959 | 0.042 |
| ○ Soil RNA vs. ✕ biofilm DNA | 0.773529 | 0.015 |

**Supplementary Figure 1.** (**A**) Acid sulfate soil taxonomic structures dissimilarity at 16S ribosomal cDNA level, as estimated using the Jaccard index. (**B**) Taxonomic structure analysis (ANOSIM) of soil 16S ribosomal DNA, soil 16S ribosomal cDNA, biofilm 16S ribosomal DNA, and biofilm 16S ribosomal cDNA.

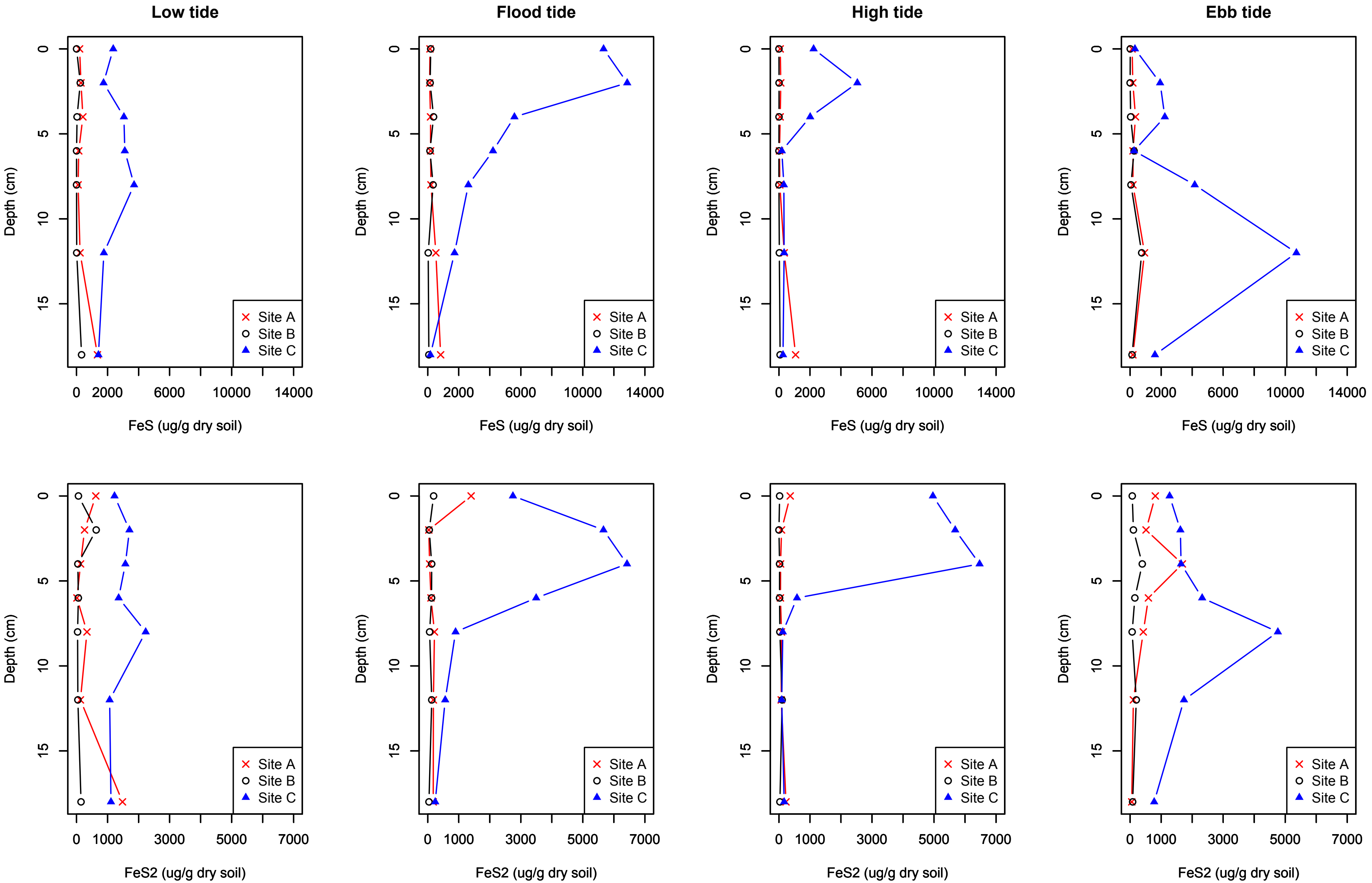


**Supplementary Figure 2.** Chemical profiles of iron monosulfide (FeS) and pyrite ($FeS_2$) in three sampling sites across four tidal stages.

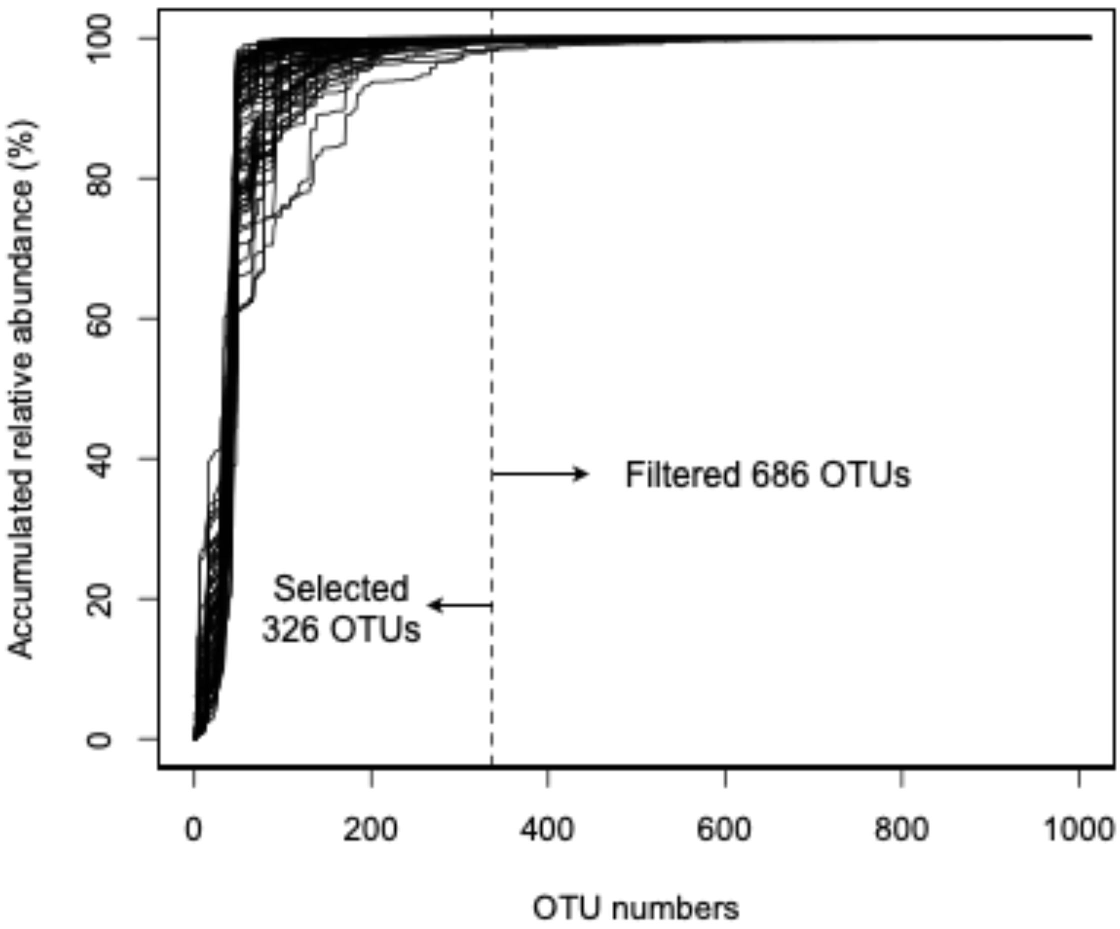


**Supplementary Figure 3.** The accumulated relative abundance of OTUs from 84 cDNA samples that were selected for training the AI package to find the best-fit pyrite predicting equation and verifying the results.

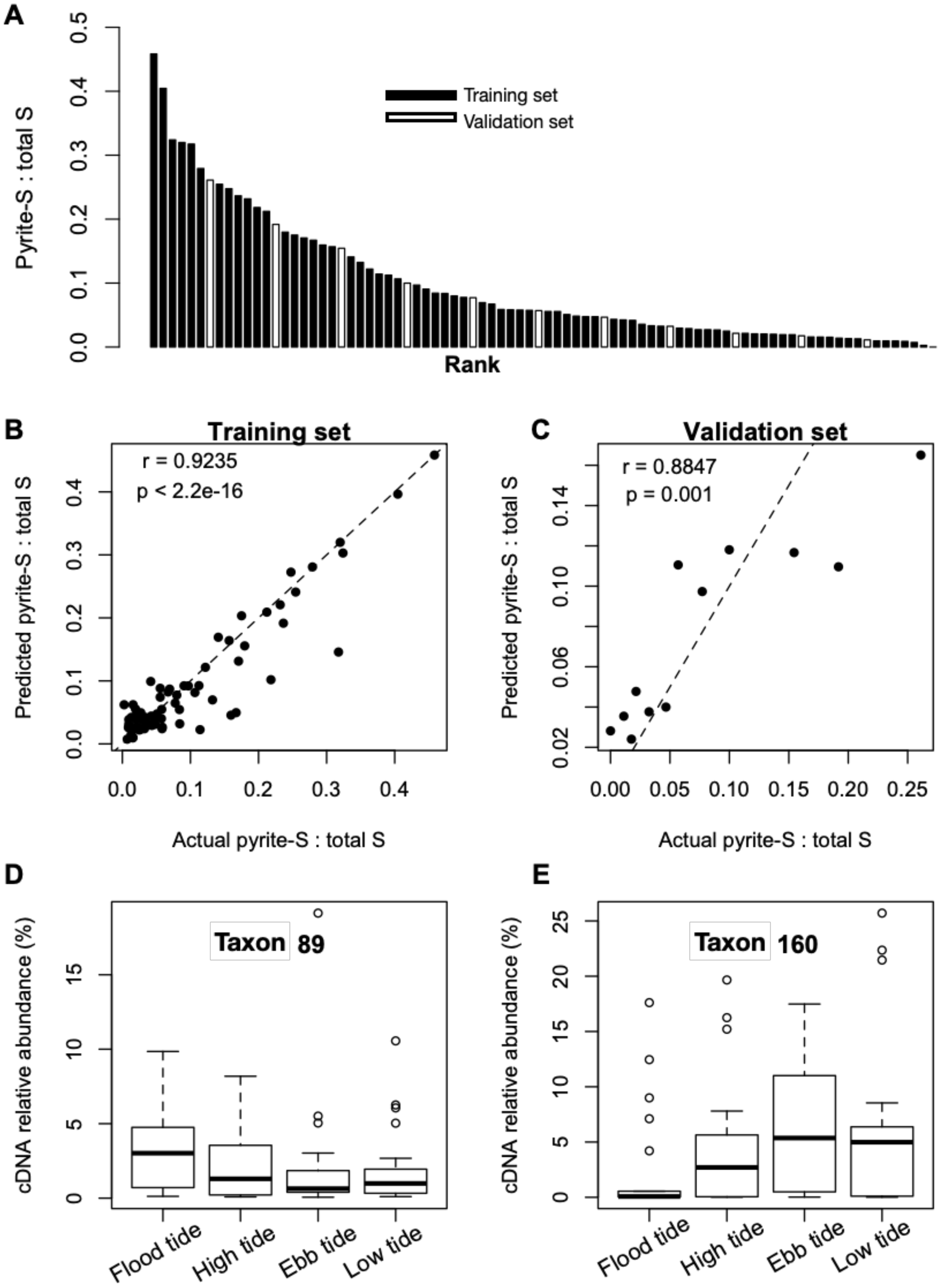


**Supplementary Figure 4.** (**A**) Total of 84 "training set" and "validation set" samples re-ranked based on the pyrite-S ratio. The training set was used to train the artificial intelligence powered modeling package to determine the best-fit equation of pyrite-S ratio, with the validation set selected to test for verification of results. (**B**, **C**) Correlations between actual and predicted pyrite-S fractions from training and validation sets, respectively. (**D**, **E**) The activity (cDNA based) of taxon89 and taxon160 across a wetting/drying cycle; these two taxa were selected using an artificial intelligence powered modeling package.

**A**

**B**

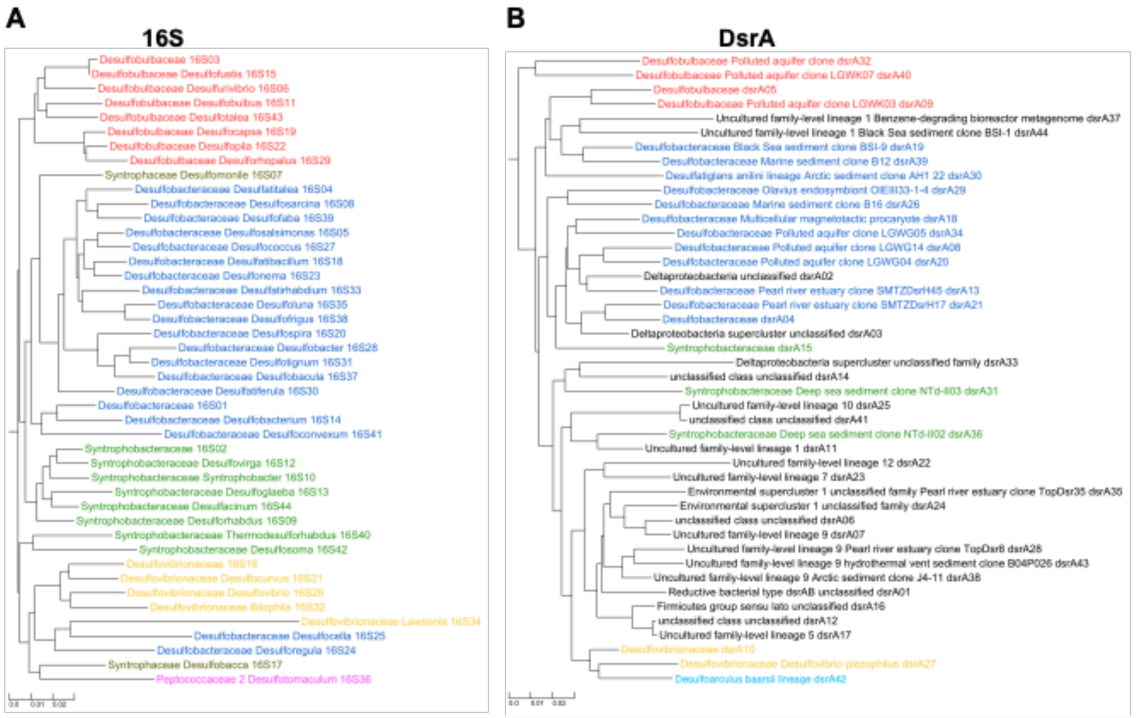


**C**

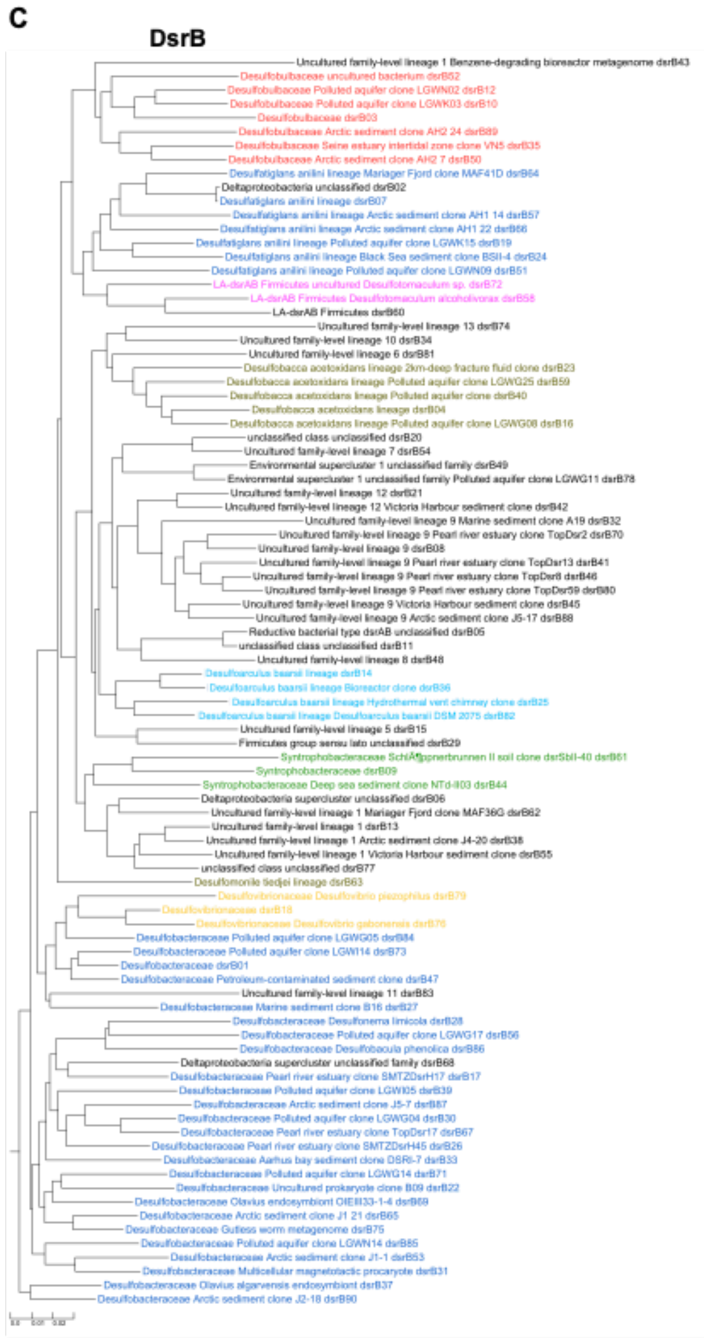


**D**

| Color | *dsrA* | *dsrB* | 16S | Kingdom | Phylum | Class | Order | Family | Genus |
|---|---|---|---|---|---|---|---|---|---|
| | x | x | x | Bacteria | Proteobacteria | Deltaproteobacteria | Desulfobacterales | Desulfobacteraceae | |
| | x | x | x | | | | | Desulfobulbaceae | |
| | x | x | | | | | | Desulfoarculaceae | Desulfoarculus |
| | x | x | x | | | | Desulfovibrionales | Desulfovibrionaceae | |
| | x | x | x | | | | Syntrophobacterales | Syntrophobacteraceae | |
| | | x | x | | | | | Syntrophaceae | Desulfobacca |
| | | x | x | | Firmicutes | Clostridia | Clostridiales | Peptococcaceae 2 | Desulfotomaculum |

**E**

| *dsrA* | *dsrB* | Level 1 | Level 2 | Level 3 | Level 4 | Level 5 |
|---|---|---|---|---|---|---|
| x | x | Reductive bacterial type dsrAB | Deltaproteobacteria supercluster | unclassified phylum | unclassified class | Uncultured family-level lineage 1 |
| | x | | | | | Uncultured family-level lineage 11 |
| x | x | | Environmental supercluster 1 | unclassified phylum | unclassified class | Uncultured family-level lineage 12 |
| | x | | | | | Uncultured family-level lineage 8 |
| x | x | | | | | Uncultured family-level lineage 9 |
| x | x | | Firmicutes group sensu lato | unclassified phylum | unclassified class | Uncultured family-level lineage 5 |
| | x | | | | | Uncultured family-level lineage 6 |
| x | x | | | | | Uncultured family-level lineage 7 |
| x | x | | Nitrospirae supercluster | unclassified phylum | unclassified class | Uncultured family-level lineage 10 |
| | x | | | | | Uncultured family-level lineage 13 |

**Supplementary Figure 5.** Consensus phylogeny of sulfate-reducing prokaryotes as identified by the following gene sequences (**A**) 16S rRNA, (**B**) *dsrA*, and (**C**) *dsrB*. (**D**) Taxonomic classification of SRP based on *dsrA* and *dsrB* sequences. (**E**) The taxonomic classification of uncultured SRP based on *dsrA* and *dsrB* sequences.

### 2. Supplementary Tables

**Supplementary Table 1.** Field area locations and tidal stages from which samples were selected for analyses of environmental Fe and S speciation, microbial diversity, SRP activity, SRP diversity, and experimental reproducibility.

| | **Sites** | **Tidal stages** | **Depth** | **Triplicate subsamples** | **Sample number** |
|---|---|---|---|---|---|
| **Microbial activity (cDNA based)** | A (supra tidal zone)<br>B (inter tidal zone)<br>C (sub-tidal zone) | T1 (ebb tide)<br>T2 (low tide)<br>T3 (flood tide)<br>T4 (high tide) | 1 (0-2 cm)<br>2 (2-4 cm)<br>3 (4-6 cm)<br>4 (6-8 cm)<br>5 (8-10 cm)<br>6 (12-14 cm)<br>7 (18-20 cm) | Homogenised | 84 |
| **Environmental Fe and S species** | A, B, C | T1, T2, T3, T4 | 1, 2, 3, 4, 5, 6, 7 | Homogenised | 84 |
| **Microbial diversity (DNA based)** | A, B, C | Homogenised | 1, 2, 3, 4, 5, 6, 7 | Homogenised | 21 |
| **SRP activity (cDNA based *dsrA* and *dsrB*)** | A, B, C | T1, T2, T3, T4 | Homogenised | Homogenised | 12 |
| **SRP diversity (DNA based *dsrA* and *dsrB*)** | A, B, C | Homogenised | Homogenised | Homogenised | 3 |
| **Reproducibility (DNA based)** | A, B, C | T1, T2, T3, T4 | 2, 4, 6 | Select 2 of them | 72 |
| **Dissimilarity between DNA and RNA** | Biofilm | T4 | Surfacewater | NA | 2 |

**Supplementary Table 2.** Chemical extractants used to differentiate operationally defined Fe or S fractions in acid sulfate soil samples, and ratios used to convert fractions to "iron (Fe) factors".

| Extractant | Fraction defined | Converted to Fe factor |
|---|---|---|
| $MgCl_2$-Fe | Soluble Fe salts | $MgCl_2$-Fe |
| HCl-Fe | Poorly crystalline Fe (after minus [FeS-Fe]) | HCl-Fe |
| CDB-Fe | Crystalline Fe | CDB-Fe |
| AVS-S | S in FeS | [FeS-S] : [FeS-Fe] = 1 : 1 |
| ES-S | Elemental sulfur | |
| CRS-S | S in $FeS_2$ | [$FeS_2$-S] : [$FeS_2$-Fe] = 2 : 1 |

**Supplementary Table 3.** OTUs within taxa in Table 1

| **Taxon** | **taxon089** | | | | | |
|---|---|---|---|---|---|---|
| **OTU** | **Kingdom** | **Phylum** | **Class** | **Order** | **Family** | **Genus** |
| OTU068 | Bacteria | Acidobacteria | Acidobacteria Gp3 | Candidatus_Solibacter | unclassified | unclassified |
| OTU069 | | | | Gp3 | unclassified | unclassified |
| OTU070 | | | | unclassified | unclassified | unclassified |

| **Taxon** | **taxon492** | | | | | |
|---|---|---|---|---|---|---|
| **OTU** | **Kingdom** | **Phylum** | **Class** | **Order** | **Family** | **Genus** |
| OTU125 | Bacteria | Actinobacteria | Actinobacteria | Actinomycetales | Microbacteriaceae | Amnibacterium |
| OTU126 | | | | | | Curtobacterium |

| **Taxon** | **taxon694** | | | | | |
|---|---|---|---|---|---|---|
| **OTU** | **Kingdom** | **Phylum** | **Class** | **Order** | **Family** | **Genus** |
| OTU647 | Bacteria | Proteobacteria | Alphaproteobacteria | Rhizobiales | Methylobacteriaceae | Methylobacterium |
| OTU648 | | | | | | Microvirga |

| **Taxon** | **taxon160** | | | | | |
|---|---|---|---|---|---|---|
| **OTU** | **Kingdom** | **Phylum** | **Class** | **Order** | **Family** | **Genus** |
| OTU0956 | Bacteria | Proteobacteria | Gammaproteobacteria | Acidithiobacillales | unclassified | unclassified |
| OTU0957 | Bacteria | Proteobacteria | Gammaproteobacteria | Aeromonadales | Aeromonadaceae | Aeromonas |
| OTU0958 | Bacteria | Proteobacteria | Gammaproteobacteria | Aeromonadales | Aeromonadaceae | Oceanimonas |
| OTU0959 | Bacteria | Proteobacteria | Gammaproteobacteria | Aeromonadales | Succinivibrionaceae | Succinivibrio |
| OTU0960 | Bacteria | Proteobacteria | Gammaproteobacteria | Alteromonadales | Alteromonadaceae | Aestuariibacter |
| OTU0961 | Bacteria | Proteobacteria | Gammaproteobacteria | Alteromonadales | Alteromonadaceae | Alishewanella |
| OTU0962 | Bacteria | Proteobacteria | Gammaproteobacteria | Alteromonadales | Alteromonadaceae | Alteromonas |
| OTU0963 | Bacteria | Proteobacteria | Gammaproteobacteria | Alteromonadales | Alteromonadaceae | Bowmanella |
| OTU0964 | Bacteria | Proteobacteria | Gammaproteobacteria | Alteromonadales | Alteromonadaceae | Glaciecola |
| OTU0965 | Bacteria | Proteobacteria | Gammaproteobacteria | Alteromonadales | Alteromonadaceae | Haliea |
| OTU0966 | Bacteria | Proteobacteria | Gammaproteobacteria | Alteromonadales | Alteromonadaceae | Marinobacter |
| OTU0967 | Bacteria | Proteobacteria | Gammaproteobacteria | Alteromonadales | Alteromonadaceae | Marinobacterium |

| | | | | | | |
|---|---|---|---|---|---|---|
| OTU0968 | Bacteria | Proteobacteria | Gammaproteobacteria | Alteromonadales | Alteromonadaceae | Microbulbifer |
| OTU0969 | Bacteria | Proteobacteria | Gammaproteobacteria | Alteromonadales | Alteromonadaceae | Salinimonas |
| OTU0970 | Bacteria | Proteobacteria | Gammaproteobacteria | Alteromonadales | Alteromonadaceae | unclassified |
| OTU0971 | Bacteria | Proteobacteria | Gammaproteobacteria | Alteromonadales | Alteromonadales_incertae_sedis | Teredinibacter |
| OTU0972 | Bacteria | Proteobacteria | Gammaproteobacteria | Alteromonadales | Celerinatantimonadaceae | Celerinatantimonas |
| OTU0973 | Bacteria | Proteobacteria | Gammaproteobacteria | Alteromonadales | Colwelliaceae | Thalassomonas |
| OTU0974 | Bacteria | Proteobacteria | Gammaproteobacteria | Alteromonadales | Colwelliaceae | unclassified |
| OTU0975 | Bacteria | Proteobacteria | Gammaproteobacteria | Alteromonadales | Ferrimonadaceae | Ferrimonas |
| OTU0976 | Bacteria | Proteobacteria | Gammaproteobacteria | Alteromonadales | Ferrimonadaceae | unclassified |
| OTU0977 | Bacteria | Proteobacteria | Gammaproteobacteria | Alteromonadales | Idiomarinaceae | Idiomarina |
| OTU0978 | Bacteria | Proteobacteria | Gammaproteobacteria | Alteromonadales | Idiomarinaceae | unclassified |
| OTU0979 | Bacteria | Proteobacteria | Gammaproteobacteria | Alteromonadales | Maricurvus | unclassified |
| OTU0980 | Bacteria | Proteobacteria | Gammaproteobacteria | Alteromonadales | Moritellaceae | Moritella |
| OTU0981 | Bacteria | Proteobacteria | Gammaproteobacteria | Alteromonadales | Pseudoalteromonadaceae | Pseudoalteromonas |
| OTU0982 | Bacteria | Proteobacteria | Gammaproteobacteria | Alteromonadales | Pseudoalteromonadaceae | Psychrosphaera |
| OTU0983 | Bacteria | Proteobacteria | Gammaproteobacteria | Alteromonadales | Pseudoalteromonadaceae | unclassified |
| OTU0984 | Bacteria | Proteobacteria | Gammaproteobacteria | Alteromonadales | Psychromonadaceae | Psychromonas |
| OTU0985 | Bacteria | Proteobacteria | Gammaproteobacteria | Alteromonadales | Shewanellaceae | Shewanella |
| OTU0986 | Bacteria | Proteobacteria | Gammaproteobacteria | Alteromonadales | unclassified | unclassified |
| OTU0987 | Bacteria | Proteobacteria | Gammaproteobacteria | Cardiobacteriales | Cardiobacteriaceae | Cardiobacterium |
| OTU0988 | Bacteria | Proteobacteria | Gammaproteobacteria | Chromatiales | Chromatiaceae | Halochromatium |
| OTU0989 | Bacteria | Proteobacteria | Gammaproteobacteria | Chromatiales | Chromatiaceae | Marichromatium |
| OTU0990 | Bacteria | Proteobacteria | Gammaproteobacteria | Chromatiales | Chromatiaceae | Rheinheimera |
| OTU0991 | Bacteria | Proteobacteria | Gammaproteobacteria | Chromatiales | Chromatiaceae | Thiohalocapsa |
| OTU0992 | Bacteria | Proteobacteria | Gammaproteobacteria | Chromatiales | Chromatiaceae | Thiophaeococcus |
| OTU0993 | Bacteria | Proteobacteria | Gammaproteobacteria | Chromatiales | Chromatiaceae | Thiorhodococcus |
| OTU0994 | Bacteria | Proteobacteria | Gammaproteobacteria | Chromatiales | Chromatiaceae | unclassified |
| OTU0995 | Bacteria | Proteobacteria | Gammaproteobacteria | Chromatiales | Ectothiorhodospiraceae | Alkalilimnicola |
| OTU0996 | Bacteria | Proteobacteria | Gammaproteobacteria | Chromatiales | Ectothiorhodospiraceae | Alkalispirillum |
| OTU0997 | Bacteria | Proteobacteria | Gammaproteobacteria | Chromatiales | Ectothiorhodospiraceae | Arhodomonas |
| OTU0998 | Bacteria | Proteobacteria | Gammaproteobacteria | Chromatiales | Ectothiorhodospiraceae | Nitrococcus |
| OTU0999 | Bacteria | Proteobacteria | Gammaproteobacteria | Chromatiales | Ectothiorhodospiraceae | Thioalbus |
| OTU1000 | Bacteria | Proteobacteria | Gammaproteobacteria | Chromatiales | Ectothiorhodospiraceae | Thioalkalispira |
| OTU1001 | Bacteria | Proteobacteria | Gammaproteobacteria | Chromatiales | Ectothiorhodospiraceae | Thioalkalivibrio |
| OTU1002 | Bacteria | Proteobacteria | Gammaproteobacteria | Chromatiales | Ectothiorhodospiraceae | Thiohalospira |
| OTU1003 | Bacteria | Proteobacteria | Gammaproteobacteria | Chromatiales | Ectothiorhodospiraceae | unclassified |
| OTU1004 | Bacteria | Proteobacteria | Gammaproteobacteria | Chromatiales | Granulosicoccaceae | Granulosicoccus |
| OTU1005 | Bacteria | Proteobacteria | Gammaproteobacteria | Chromatiales | Halothiobacillaceae | Halothiobacillus |
| OTU1006 | Bacteria | Proteobacteria | Gammaproteobacteria | Chromatiales | Halothiobacillaceae | Thioalkalibacter |
| OTU1007 | Bacteria | Proteobacteria | Gammaproteobacteria | Chromatiales | Halothiobacillaceae | unclassified |
| OTU1008 | Bacteria | Proteobacteria | Gammaproteobacteria | Chromatiales | unclassified | unclassified |
| OTU1009 | Bacteria | Proteobacteria | Gammaproteobacteria | Enterobacteriales | Enterobacteriaceae | Buttiauxella |
| OTU1010 | Bacteria | Proteobacteria | Gammaproteobacteria | Enterobacteriales | Enterobacteriaceae | Cronobacter |
| OTU1011 | Bacteria | Proteobacteria | Gammaproteobacteria | Enterobacteriales | Enterobacteriaceae | Enterobacter |
| OTU1012 | Bacteria | Proteobacteria | Gammaproteobacteria | Enterobacteriales | Enterobacteriaceae | Escherichia/Shigella |
| OTU1013 | Bacteria | Proteobacteria | Gammaproteobacteria | Enterobacteriales | Enterobacteriaceae | Klebsiella |

| | | | | | | |
|---|---|---|---|---|---|---|
| OTU1014 | Bacteria | Proteobacteria | Gammaproteobacteria | Enterobacteriales | Enterobacteriaceae | Morganella |
| OTU1015 | Bacteria | Proteobacteria | Gammaproteobacteria | Enterobacteriales | Enterobacteriaceae | Pantoea |
| OTU1016 | Bacteria | Proteobacteria | Gammaproteobacteria | Enterobacteriales | Enterobacteriaceae | Salmonella |
| OTU1017 | Bacteria | Proteobacteria | Gammaproteobacteria | Enterobacteriales | Enterobacteriaceae | Serratia |
| OTU1018 | Bacteria | Proteobacteria | Gammaproteobacteria | Enterobacteriales | Enterobacteriaceae | unclassified |
| OTU1019 | Bacteria | Proteobacteria | Gammaproteobacteria | Gammaproteobacteria_ incertae_sedis | Arenicella | unclassified |
| OTU1020 | Bacteria | Proteobacteria | Gammaproteobacteria | Gammaproteobacteria_ incertae_sedis | Congregibacter | unclassified |
| OTU1021 | Bacteria | Proteobacteria | Gammaproteobacteria | Gammaproteobacteria_ incertae_sedis | Eionea | unclassified |
| OTU1022 | Bacteria | Proteobacteria | Gammaproteobacteria | Gammaproteobacteria_ incertae_sedis | Gallaecimonas | unclassified |
| OTU1023 | Bacteria | Proteobacteria | Gammaproteobacteria | Gammaproteobacteria_ incertae_sedis | Gilvimarinus | unclassified |
| OTU1024 | Bacteria | Proteobacteria | Gammaproteobacteria | Gammaproteobacteria_ incertae_sedis | Marinicella | unclassified |
| OTU1025 | Bacteria | Proteobacteria | Gammaproteobacteria | Gammaproteobacteria_ incertae_sedis | Methylohalomonas | unclassified |
| OTU1026 | Bacteria | Proteobacteria | Gammaproteobacteria | Gammaproteobacteria_ incertae_sedis | Methylonatrum | unclassified |
| OTU1027 | Bacteria | Proteobacteria | Gammaproteobacteria | Gammaproteobacteria_ incertae_sedis | Porticoccus | unclassified |
| OTU1028 | Bacteria | Proteobacteria | Gammaproteobacteria | Gammaproteobacteria_ incertae_sedis | Sedimenticola | unclassified |
| OTU1029 | Bacteria | Proteobacteria | Gammaproteobacteria | Gammaproteobacteria_ incertae_sedis | Solimonas | unclassified |
| OTU1030 | Bacteria | Proteobacteria | Gammaproteobacteria | Gammaproteobacteria_ incertae_sedis | Spongiibacter | unclassified |
| OTU1031 | Bacteria | Proteobacteria | Gammaproteobacteria | Gammaproteobacteria_ incertae_sedis | Thiohalobacter | unclassified |
| OTU1032 | Bacteria | Proteobacteria | Gammaproteobacteria | Gammaproteobacteria_ incertae_sedis | Thiohalomonas | unclassified |
| OTU1033 | Bacteria | Proteobacteria | Gammaproteobacteria | Gammaproteobacteria_ incertae_sedis | Thiohalophilus | unclassified |
| OTU1034 | Bacteria | Proteobacteria | Gammaproteobacteria | Gammaproteobacteria_ incertae_sedis | Thioprofundum | unclassified |
| OTU1035 | Bacteria | Proteobacteria | Gammaproteobacteria | Gammaproteobacteria_ incertae_sedis | unclassified | unclassified |
| OTU1036 | Bacteria | Proteobacteria | Gammaproteobacteria | Halioglobus | unclassified | unclassified |
| OTU1037 | Bacteria | Proteobacteria | Gammaproteobacteria | Legionellales | Coxiellaceae | Aquicella |
| OTU1038 | Bacteria | Proteobacteria | Gammaproteobacteria | Legionellales | Coxiellaceae | Coxiella |
| OTU1039 | Bacteria | Proteobacteria | Gammaproteobacteria | Legionellales | Coxiellaceae | Diplorickettsia |
| OTU1040 | Bacteria | Proteobacteria | Gammaproteobacteria | Legionellales | Coxiellaceae | unclassified |
| OTU1041 | Bacteria | Proteobacteria | Gammaproteobacteria | Legionellales | Legionellaceae | Legionella |
| OTU1042 | Bacteria | Proteobacteria | Gammaproteobacteria | Methylococcales | Methylococcaceae | Methylobacter |
| OTU1043 | Bacteria | Proteobacteria | Gammaproteobacteria | Methylococcales | Methylococcaceae | Methylococcus |
| OTU1044 | Bacteria | Proteobacteria | Gammaproteobacteria | Methylococcales | Methylococcaceae | Methylomarinum |
| OTU1045 | Bacteria | Proteobacteria | Gammaproteobacteria | Methylococcales | Methylococcaceae | Methylomicrobium |
| OTU1046 | Bacteria | Proteobacteria | Gammaproteobacteria | Methylococcales | Methylococcaceae | Methylomonas |
| OTU1047 | Bacteria | Proteobacteria | Gammaproteobacteria | Methylococcales | Methylococcaceae | Methylosarcina |
| OTU1048 | Bacteria | Proteobacteria | Gammaproteobacteria | Methylococcales | Methylococcaceae | unclassified |
| OTU1049 | Bacteria | Proteobacteria | Gammaproteobacteria | Oceanospirillales | Alcanivoracaceae | Alcanivorax |
| OTU1050 | Bacteria | Proteobacteria | Gammaproteobacteria | Oceanospirillales | Hahellaceae | Endozoicomonas |
| OTU1051 | Bacteria | Proteobacteria | Gammaproteobacteria | Oceanospirillales | Hahellaceae | Hahella |
| OTU1052 | Bacteria | Proteobacteria | Gammaproteobacteria | Oceanospirillales | Hahellaceae | Zooshikella |
| OTU1053 | Bacteria | Proteobacteria | Gammaproteobacteria | Oceanospirillales | Halomonadaceae | Halomonas |
| OTU1054 | Bacteria | Proteobacteria | Gammaproteobacteria | Oceanospirillales | Halomonadaceae | Kushneria |

| | | | | | | |
|---|---|---|---|---|---|---|
| OTU1055 | Bacteria | Proteobacteria | Gammaproteobacteria | Oceanospirillales | Halomonadaceae | Modicisalibacter |
| OTU1056 | Bacteria | Proteobacteria | Gammaproteobacteria | Oceanospirillales | Halomonadaceae | Salinicola |
| OTU1057 | Bacteria | Proteobacteria | Gammaproteobacteria | Oceanospirillales | Halomonadaceae | unclassified |
| OTU1058 | Bacteria | Proteobacteria | Gammaproteobacteria | Oceanospirillales | Oceanospirillaceae | Amphritea |
| OTU1059 | Bacteria | Proteobacteria | Gammaproteobacteria | Oceanospirillales | Oceanospirillaceae | Bermanella |
| OTU1060 | Bacteria | Proteobacteria | Gammaproteobacteria | Oceanospirillales | Oceanospirillaceae | Corallomonas |
| OTU1061 | Bacteria | Proteobacteria | Gammaproteobacteria | Oceanospirillales | Oceanospirillaceae | Marinomonas |
| OTU1062 | Bacteria | Proteobacteria | Gammaproteobacteria | Oceanospirillales | Oceanospirillaceae | Neptuniibacter |
| OTU1063 | Bacteria | Proteobacteria | Gammaproteobacteria | Oceanospirillales | Oceanospirillaceae | Neptunomonas |
| OTU1064 | Bacteria | Proteobacteria | Gammaproteobacteria | Oceanospirillales | Oceanospirillaceae | Nitrincola |
| OTU1065 | Bacteria | Proteobacteria | Gammaproteobacteria | Oceanospirillales | Oceanospirillaceae | Oceaniserpentilla |
| OTU1066 | Bacteria | Proteobacteria | Gammaproteobacteria | Oceanospirillales | Oceanospirillaceae | Oceanobacter |
| OTU1067 | Bacteria | Proteobacteria | Gammaproteobacteria | Oceanospirillales | Oceanospirillaceae | Oceanospirillum |
| OTU1068 | Bacteria | Proteobacteria | Gammaproteobacteria | Oceanospirillales | Oceanospirillaceae | Reinekea |
| OTU1069 | Bacteria | Proteobacteria | Gammaproteobacteria | Oceanospirillales | Oceanospirillaceae | Thalassolituus |
| OTU1070 | Bacteria | Proteobacteria | Gammaproteobacteria | Oceanospirillales | Oceanospirillaceae | unclassified |
| OTU1071 | Bacteria | Proteobacteria | Gammaproteobacteria | Oceanospirillales | Oleiphilaceae | Oleiphilus |
| OTU1072 | Bacteria | Proteobacteria | Gammaproteobacteria | Oceanospirillales | Saccharospirillaceae | Saccharospirillum |
| OTU1073 | Bacteria | Proteobacteria | Gammaproteobacteria | Oceanospirillales | unclassified | unclassified |
| OTU1074 | Bacteria | Proteobacteria | Gammaproteobacteria | Orbales | Orbaceae | unclassified |
| OTU1075 | Bacteria | Proteobacteria | Gammaproteobacteria | Pasteurellales | Pasteurellaceae | Aggregatibacter |
| OTU1076 | Bacteria | Proteobacteria | Gammaproteobacteria | Pasteurellales | Pasteurellaceae | Haemophilus |
| OTU1077 | Bacteria | Proteobacteria | Gammaproteobacteria | Pasteurellales | Pasteurellaceae | Mannheimia |
| OTU1078 | Bacteria | Proteobacteria | Gammaproteobacteria | Pasteurellales | Pasteurellaceae | unclassified |
| OTU1079 | Bacteria | Proteobacteria | Gammaproteobacteria | Pseudomonadales | Moraxellaceae | Acinetobacter |
| OTU1080 | Bacteria | Proteobacteria | Gammaproteobacteria | Pseudomonadales | Moraxellaceae | Enhydrobacter |
| OTU1081 | Bacteria | Proteobacteria | Gammaproteobacteria | Pseudomonadales | Moraxellaceae | Psychrobacter |
| OTU1082 | Bacteria | Proteobacteria | Gammaproteobacteria | Pseudomonadales | Moraxellaceae | unclassified |
| OTU1083 | Bacteria | Proteobacteria | Gammaproteobacteria | Pseudomonadales | Pseudomonadaceae | Cellvibrio |
| OTU1084 | Bacteria | Proteobacteria | Gammaproteobacteria | Pseudomonadales | Pseudomonadaceae | Pseudomonas |
| OTU1085 | Bacteria | Proteobacteria | Gammaproteobacteria | Pseudomonadales | Pseudomonadaceae | unclassified |
| OTU1086 | Bacteria | Proteobacteria | Gammaproteobacteria | Pseudomonadales | Pseudomonadales_incertae_sedis | Dasania |
| OTU1087 | Bacteria | Proteobacteria | Gammaproteobacteria | Pseudomonadales | unclassified | unclassified |
| OTU1088 | Bacteria | Proteobacteria | Gammaproteobacteria | Salinisphaerales | Salinisphaeraceae | Salinisphaera |
| OTU1089 | Bacteria | Proteobacteria | Gammaproteobacteria | Thiotrichales | Francisellaceae | Francisella |
| OTU1090 | Bacteria | Proteobacteria | Gammaproteobacteria | Thiotrichales | Piscirickettsiaceae | Hydrogenovibrio |
| OTU1091 | Bacteria | Proteobacteria | Gammaproteobacteria | Thiotrichales | Piscirickettsiaceae | Methylophaga |
| OTU1092 | Bacteria | Proteobacteria | Gammaproteobacteria | Thiotrichales | Piscirickettsiaceae | Thiomicrospira |
| OTU1093 | Bacteria | Proteobacteria | Gammaproteobacteria | Thiotrichales | Piscirickettsiaceae | unclassified |
| OTU1094 | Bacteria | Proteobacteria | Gammaproteobacteria | Thiotrichales | Thiotrichaceae | Thiothrix |
| OTU1095 | Bacteria | Proteobacteria | Gammaproteobacteria | Thiotrichales | Thiotrichales_incertae_sedis | Fangia |
| OTU1096 | Bacteria | Proteobacteria | Gammaproteobacteria | Thiotrichales | unclassified | unclassified |
| OTU1097 | Bacteria | Proteobacteria | Gammaproteobacteria | Vibrionales | Vibrionaceae | Allomonas |
| OTU1098 | Bacteria | Proteobacteria | Gammaproteobacteria | Vibrionales | Vibrionaceae | Enterovibrio |
| OTU1099 | Bacteria | Proteobacteria | Gammaproteobacteria | Vibrionales | Vibrionaceae | Grimontia |
| OTU1100 | Bacteria | Proteobacteria | Gammaproteobacteria | Vibrionales | Vibrionaceae | Lucibacterium |

| | | | | | | |
|---|---|---|---|---|---|---|
| OTU1101 | Bacteria | Proteobacteria | Gammaproteobacteria | Vibrionales | Vibrionaceae | Photobacterium |
| OTU1102 | Bacteria | Proteobacteria | Gammaproteobacteria | Vibrionales | Vibrionaceae | Salinivibrio |
| OTU1103 | Bacteria | Proteobacteria | Gammaproteobacteria | Vibrionales | Vibrionaceae | Vibrio |
| OTU1104 | Bacteria | Proteobacteria | Gammaproteobacteria | Vibrionales | Vibrionaceae | unclassified |
| OTU1105 | Bacteria | Proteobacteria | Gammaproteobacteria | Xanthomonadales | Sinobacteraceae | Alkanibacter |
| OTU1106 | Bacteria | Proteobacteria | Gammaproteobacteria | Xanthomonadales | Sinobacteraceae | Nevskia |
| OTU1107 | Bacteria | Proteobacteria | Gammaproteobacteria | Xanthomonadales | Sinobacteraceae | Singularimonas |
| OTU1108 | Bacteria | Proteobacteria | Gammaproteobacteria | Xanthomonadales | Sinobacteraceae | Steroidobacter |
| OTU1109 | Bacteria | Proteobacteria | Gammaproteobacteria | Xanthomonadales | Sinobacteraceae | unclassified |
| OTU1110 | Bacteria | Proteobacteria | Gammaproteobacteria | Xanthomonadales | Xanthomonadaceae | Dokdonella |
| OTU1111 | Bacteria | Proteobacteria | Gammaproteobacteria | Xanthomonadales | Xanthomonadaceae | Dyella |
| OTU1112 | Bacteria | Proteobacteria | Gammaproteobacteria | Xanthomonadales | Xanthomonadaceae | Luteibacter |
| OTU1113 | Bacteria | Proteobacteria | Gammaproteobacteria | Xanthomonadales | Xanthomonadaceae | Lysobacter |
| OTU1114 | Bacteria | Proteobacteria | Gammaproteobacteria | Xanthomonadales | Xanthomonadaceae | Pseudoxanthomonas |
| OTU1115 | Bacteria | Proteobacteria | Gammaproteobacteria | Xanthomonadales | Xanthomonadaceae | Rudaea |
| OTU1116 | Bacteria | Proteobacteria | Gammaproteobacteria | Xanthomonadales | Xanthomonadaceae | Stenotrophomonas |
| OTU1117 | Bacteria | Proteobacteria | Gammaproteobacteria | Xanthomonadales | Xanthomonadaceae | Xanthomonas |
| OTU1118 | Bacteria | Proteobacteria | Gammaproteobacteria | Xanthomonadales | Xanthomonadaceae | unclassified |
| OTU1119 | Bacteria | Proteobacteria | Gammaproteobacteria | Xanthomonadales | unclassified | unclassified |
| OTU1120 | Bacteria | Proteobacteria | Gammaproteobacteria | unclassified | unclassified | unclassified |
